# SELEX-HTCFQ Platform: Developing DNA Enhancers of ADAR1 to Suppress ZBP1-Dependent Immunopathology

**DOI:** 10.1101/2025.10.22.683608

**Authors:** Yusi Hu, Rentang Huang, Yi-Fan Wang, Han Zhu, Qing-Qing Ye, Juan-Mei Wang, Zhuhua Yao, Zhi-Gang Wang, Dai-Wen Pang, Shu-Lin Liu

**Affiliations:** International Joint Research Center of Human-machine Intelligent, Collaborative for Tumor Precision Diagnosis and Treatment of Hainan Province & Key Laboratory of Tropical Translational Medicine of Ministry of Education, School of Pharmacy & The First Affiliated Hospital, Hainan Medical University, Haikou 571199, P. R. China; State Key Laboratory of Medicinal Chemical Biology, Frontiers Science Centre for New Organic Matter, Tianjin Key Laboratory of Biosensing and Molecular Recognition, Research Centre for Analytical Sciences, College of Chemistry, School of Medicine, and Frontiers Science Centre for Cell Responses, Nankai University, Tianjin 300071, P. R. China; School of Medicine, Tianjin Union Medical Center and Frontiers Science Center for Cell Responses, Nankai University, Tianjin 300071, P. R. China

**Keywords:** ADAR1, enhancer, ZBP1, DNA aptamer, inflammation

## Abstract

Excessive immune activation drives pathological inflammation through dysregulated ZBP1 signaling, yet this sensor remains crucial for immune surveillance, necessitating targeted therapies that selectively inhibit pathology while preserving protective functions. Here, we developed an innovative SELEX-HTCFQ platform that combines Systematic Evolution of Ligands by Exponential Enrichment (SELEX) with high-throughput competitive fluorescence quenching (HTCFQ) to identify ADAR1-specific enhancers. Capitalizing on ADAR1’s natural ability to suppress ZBP1 *via* competitive Z-nucleic acid binding, this platform’s dual assessment of affinity and selectivity identified aptamer A4, a highly specific ADAR1 enhancers demonstrating over 40-fold selectivity over ZBP1. A4 allosterically modulates ADAR1’s activity, thereby potentiating its inhibitory effect on ZBP1, which not only mitigates the excessive inflammatory response but also maintains the delicate balance of the immune system. These ADAR1 enhancers represent precision molecular tools for reprogramming Z-nucleic acid sensing, offering a promising therapeutic paradigm for cytokine storm syndromes and necroptosis-driven disorders through selective pathway modulation.

## Introduction

Excessive immune activation leading to pathological tissue damage represents a fundamental challenge in treating inflammatory diseases.^1, 2^ Central to this process is ZBP1 (Z-DNA binding protein 1), an innate immune sensor that recognizes viral and endogenous Z-form nucleic acids (Z-RNA/DNA) through its Zα domain.^3-5^ Upon activation, ZBP1 initiates a self-amplifying cycle of inflammation and programmed cell death by recruiting RIPK1 and RIPK3 via its RHIM domain to form the PANoptosome - a multiprotein complex that coordinately activates necroptosis (through RIPK3-MLKL signaling) and pyroptosis (via NLRP3 inflammasome-caspase-1 activation).^6-8^ This dual cell death pathway triggers massive release of pro-inflammatory cytokines (IL-1β, IL-18) and damage-associated molecular patterns (DAMPs like HMGB1), creating a feedforward loop of inflammation that disrupts tissue barriers and causes organ dysfunction.^3, 6^ The pathological significance of ZBP1 is evident in diverse conditions including viral pneumonia, chemotherapy-induced intestinal damage, and inflammatory bowel disease, making its precise regulation a critical therapeutic target.

Current therapeutic approaches, which mainly target downstream molecules of ZBP1 activation like RIPK3 and MLKL, have limitations as they fail to comprehensively tackle the issues caused by excessive ZBP1 activation and cannot stop the inflammation development and disease progression from the root.^9^ Directly inhibiting ZBP1, located at the front of the signaling pathway and controlling multiple branches, offers a more efficient and precise way to modulate inflammatory responses, with less impact on other physiological pathways and thus lower potential side effects.^10^ Moreover, inhibiting ZBP1 activity early can provide preventive protection against inflammation and cell death. Our team has developed PROTACs targeting ZBP1, achieving selective degradation of overactive ZBP1 and offering new hope for ZBP1-related diseases.^11^ However, a key challenge remains: while overactive ZBP1 drives immunopathology, its basic activity is crucial for immune surveillance and homeostasis. Thus, complete degradation of ZBP1 might weaken the host’s defense against infections and tumors,^7, 12, 13^ underscoring the need for a novel strategy that precisely inhibits pathological signaling while preserving ZBP1’s physiological functions.

ADAR1 (adenosine deaminase, RNA-specific 1) primarily regulates ZBP1 by having its Zα domain compete with ZBP1 for binding to Z-nucleic acids.^14-16^ These complementary actions position ADAR1 as a molecular brake for ZBP1 activation, preventing aberrant inflammation while maintaining immune homeostasis and allowing controlled responses to genuine threats. Thus, strategically enhancing ADAR1’s binding to Z-DNA could selectively inhibit the ZBP1 pathway, offering a promising therapeutic approach for inflammatory diseases. Concurrently, DNA aptamers, as molecular interaction modulators, hold great promise. They exhibit high affinity and specificity, enabling precise targeting of various molecules and can be obtained through *in vitro* selection.^17^ DNA aptamers are easily synthesized and modified to enhance stability, prolong half-life, and improve biocompatibility.^18^ Moreover, they are robust, withstanding harsh conditions and typically non-immunogenic. Their structural plasticity allows them to precisely recognize proteins, nucleic acids, and other biomacromolecules, making them valuable tools in diagnostics and therapeutics.^19-21^ Therefore, developing DNA aptamer-based ADAR1 enhancers may effectively inhibit the activation of the ZBP1 inflammatory pathway, offering new strategies for treating related diseases.

Here, we present a screening platform termed the SELEX-HTCFQ assay, which integrates Systematic Evolution of Ligands by Exponential Enrichment (SELEX) technique with high-throughput competitive fluorescence quenching technique (HTCFQ) to efficiently identify and validate ADAR1 enhancers. Through structure-guided screening of a single-stranded DNA (ssDNA) library targeting the Zα domain of ADAR1 (ADAR1-Zα), we identified high-affinity lead sequences, which were further validated by surface plasmon resonance (SPR) analysis.^22^ Using high-throughput competitive fluorescence quenching (HTCFQ) assays with quencher-modified DNA probes against fluorescently labeled Zα domains of both ADAR1 and ZBP1 (ZBP1-Zα), we identified the optimal enhancers sequence (Figure 1a). Utilizing this strategy, we successfully developed an ADAR1 enhancer that competitively inhibit the interaction between ZBP1 and Z-nucleic acids. In an influenza virus infection model, the selected aptamers effectively suppressed the ZBP1-mediated necroptosis pathway and significantly reduced virus-induced excessive inflammatory responses. This approach effectively minimized therapeutic side effects while maintaining normal immune function, offering a promising strategy for the precise treatment of viral inflammatory diseases.

**Figure 1.**
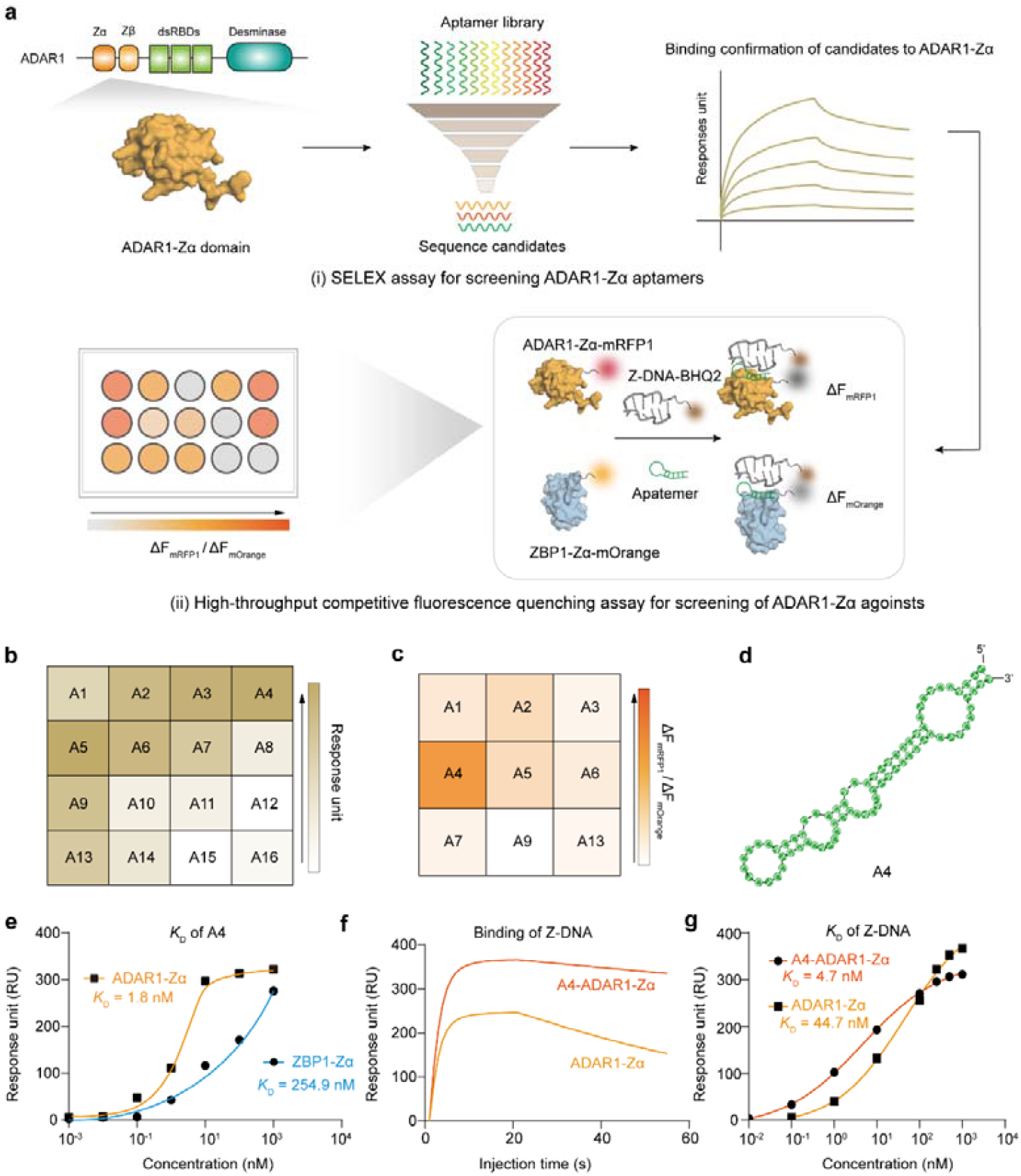
SELEX-HTCFQ screening of ADAR1-Zα enhancing aptamers. **(a)** Schematic diagram of the procedures for screening aptamer enhancers for ADAR1. Specifically, the ADAR1-Zα domain sequence was isolated from the ADAR1 sequence and expressed as a protein. Aptamers interacting with the ADAR1-Zα protein were screened using the SELEX method and analyzed by SPR (i). The top candidates were further screened using the HTCFQ assay to identify potential enhancers (ii). **(b)** SPR analysis confirming the top 16 sequences that bind most tightly to ADAR1-Zα domain. **(c)** HTCFQ identifying the top 9 sequences with high enhancing efficacy for ADAR1-Zα domain. **(d)** Predicted secondary structure of the A4 aptamer. **(e)** SPR analysis showing the binding affinity of A4 towards ADAR1-Zα (*K*_D_ = 1.8 nM) and ZBP1-Zα (*K*_D_ = 254.9 nM), respectively. **(f)** SPR binding curves of ADAR1-Zα and A4-pretreated ADAR1-Zα against Z-DNA. **(g)** SPR analysis showing the binding affinity of Z-DNA towards ADAR1 (*K*_D_ =44.7 nM) and A4-pretreated ADAR1 (*K*_D_ = 4.7 nM).

## Results and Discussion

### Screening for ADAR1 Enhancers using a SELEX-HTCFQ assay

To identify an aptamer enhancer capable of strengthening the ADAR1-Zα and Z-DNA interaction, we first aimed to select DNA aptamers that specifically bind to the ADAR1-Zα domain. Accordingly, we expressed the ADAR1-Zα (134-198) fragment in *E. coli* (Figure 1a) and subjected a large DNA library (complexity of 10^1^□, containing a 36-nt random region) to the SELEX procedure.^23-25^ After 15 rounds of selection and enrichment, high-throughput sequencing and frequency analysis identified 42 lead candidate aptamers (Table S1). From these, the 16 most frequent sequences were subjected to surface plasmon resonance (SPR) analysis to evaluate their binding affinities (Figure S1a). Subsequent SPR profiling identified the top nine binders (A1–A4, etc.), which exhibited the most robust binding to ADAR1-Zα among the candidates tested (Figure 1b). Next, to discover the most effective enhancer among these, we developed a high-throughput competitive fluorescence quenching (HTCFQ) assay leveraging fluorescence resonance energy transfer (FRET) technology. In this system, mRFP1-tagged ADAR1-Zα and mOrange-tagged ZBP1-Zα were each allowed to interact with BHQ2-labeled Z-DNA, where stronger binding resulted in greater fluorescence quenching (ΔF). By comparing the quenching efficiencies (ΔF_mRFP1_ for ADAR1-Zα versus ΔF_mOrange_ for ZBP1-Zα), we derived a selectivity ratio (ΔF_mRFP1_/ΔF_mOrange_) that quantitatively reflects each aptamer’s preferential enhancement of ADAR1-Zα-Z-DNA binding (Figure 1a). Among the nine candidates, A4 (secondary structure shown in Figure 1d) demonstrated the highest ΔF_mRFP1_/ΔF_mOrange_ value, suggesting its potential as an ADAR1-Zα enhancers (Figure 1c).

Further validation *via* SPR confirmed that the dissociation constants (*K*_D_) for A4 binding to ADAR1-Zα is 1.8 nM, while for ZBP1-Zα it is 254.9 nM, indicating that A4 binds to ADAR1-Zα with over 140 times stronger affinity than ZBP1-Zα (Figure 1e, Figure S1b-c). Additionally, pretreatment of ADAR1-Zα with A4 enhanced its binding to Z-DNA, as evidenced by higher SPR response units (Figure 1f). The affinity of A4-bound ADAR1-Zα for Z-DNA (*K*_D_ = 4.7 nM) was around 10-fold stronger than that of ADAR1-Zα alone (*K*_D_ = 44.7 nM) (Figure 1g, Figure S1d-e). Additionally, we employed the HTCFQ assay to determine the *K*_D_ of ADAR1-Zα, A4-ADAR1-Zα, and ZBP1-Zα with Z-DNA. The measured *K*_D_ values of 42.6 nM, 3.0 nM, and 70.6 nM, respectively, closely matched our SPR data (Figure S1f), reinforcing the reliability of our results. To assess competitive binding, we incubated A4-ADAR1-Zα and ZBP1-Zα together with Z-DNA. In this competitive setting, A4-ADAR1-Zα maintained strong binding (*K*_D_ = 3.5 nM), while ZBP1-Zα binding was markedly weakened (*K*_D_ = 146.7 nM) (Figure S1g). These results demonstrated that A4 selectively enhances ADAR1-Zα’s affinity for Z-DNA while competitively suppressing ZBP1-Zα binding, highlighting its potential as a specific enhancer of ADAR1-Zα activity.

### Molecular mechanism of A4-mediated enhancement of ADAR1–Z-DNA binding

To elucidate the structural basis for A4’s superior performance, we performed a multiple sequence alignment of the top 42 unique aptamers, which classified them into five distinct families (Figure S2a and Table S1). A4, the most abundant sequence in Family III, is characterized by a multi-stem-loop secondary structure. This architecture features stems rich in stable G-C base pairs and, notably, loops with a high frequency of cytosine residues (Figure S2b). To identify the functional core, we designed three truncated variants (A4-13, A4-20, and A4-32) based on this predicted structure (Figure S3a, b). SPR analysis showed that A4-13 retained binding affinity comparable to full-length A4, whereas A4-20 and A4-32 exhibited significantly weakened or completely abolished binding, respectively. These results indicated that the structural integrity of the intermediary loop regions (disrupted in A4-20 and A4-32) is critical for molecular recognition. Given the prevalence of cytosine in the loops of this core region, we hypothesized that these residues are essential for binding. To test this, we designed a series of structure-guided point mutants (A4 Loop2, A4 Loop2+3, and A4 Loop2+3+4) by substituting conserved cytosines with adenines while preserving the overall scaffold (Figure S4a, b). These mutants exhibited a stepwise decrease in binding affinity, with A4 Loop2, A4 Loop2+3, and A4 Loop2+3+4 retaining approximately 50%, 20%, and negligible levels of wild-type binding, respectively (Figure S4c). This progressive loss of function confirmed that the conserved cytosines in these loops are essential for molecular recognition.

We next investigated whether this optimized structure translates into superior binding kinetics. As a higher association rate (k_on_) enables faster target encounter and complex formation, while a lower dissociation rate (k_off_) reflects a longer-lived, more stable interaction,^26^ we performed surface plasmon resonance (SPR) analysis to evaluate these parameters. The results confirmed that A4 exhibited the second-highest association rate (k_on_) and the lowest dissociation rate (k_off_) among the top candidates, indicating that it formed the most stable complex with ADAR1-Zα (Table S2). More importantly, in the context of the ADAR1-Zα / Z-DNA interaction, A4 increased the association rate (k_on_) by approximately 1.6-fold and decreased the dissociation rate (k_off_) by about 5.6-fold. This synergistic effect resulted in a nearly 10-fold improvement in overall binding affinity (*K*_D_), reducing it from 52.3 nM to 5.7 nM (Table S3). The close agreement between kinetic (5.7 nM) and equilibrium-derived (4.7 nM) *K*_D_ values reinforced this interpretation. Together with its potent enhancer activity in an orthogonal HTCFQ assay, these results demonstrated that A4’s conserved structural features, particularly its cytosine-rich loops, were directly responsible for the enhanced binding kinetics, high affinity, and ultimate function.

Subsequently, to characterize the molecular interactions underlying A4-mediated ADAR1-Zα activation, we conducted comprehensive 200 ns molecular dynamics simulations comparing three systems: (1) Z-DNA-ADAR1-Zα, (2) A4-ADAR1-Zα, and (3) the ternary Z-DNA-A4-ADAR1-Zα complex. RMSD analysis confirmed system stability after 50 ns (Figure S5a), enabling reliable sampling of the 100-200 ns trajectory for detailed energetic and structural analyses. Free energy landscape reconstruction revealed a deep energy well for the A4-ADAR1-Zα complex (Figure S5c), demonstrating A4’s stable, low-energy binding mode that correlates with experimentally observed high affinity. Comparative hydrogen bond analysis showed A4 enhances ADAR1-Zα’s interaction with Z-DNA, increasing hydrogen bond formation from 7 (binary complex) to 12 (ternary complex) (Figures 2a-b). This structural insight provides a mechanistic explanation for A4’s enhancing effect, showing how it stabilizes ADAR1-Zα in a high-affinity Z-DNA-binding conformation. To experimentally validate these computational predictions, we performed a super-shift polyacrylamide gel electrophoresis (SPS-PAGE) assay (Figure S5b). The results showed a clear reduction in the electrophoretic mobility of the A4–ADAR1-Zα complex upon Z-DNA binding, suggesting the formation of a larger ternary complex. This observation aligns with the MD simulations, which indicated that A4 stabilizes the ADAR1-Zα–Z-DNA interface.

**Figure 2.**
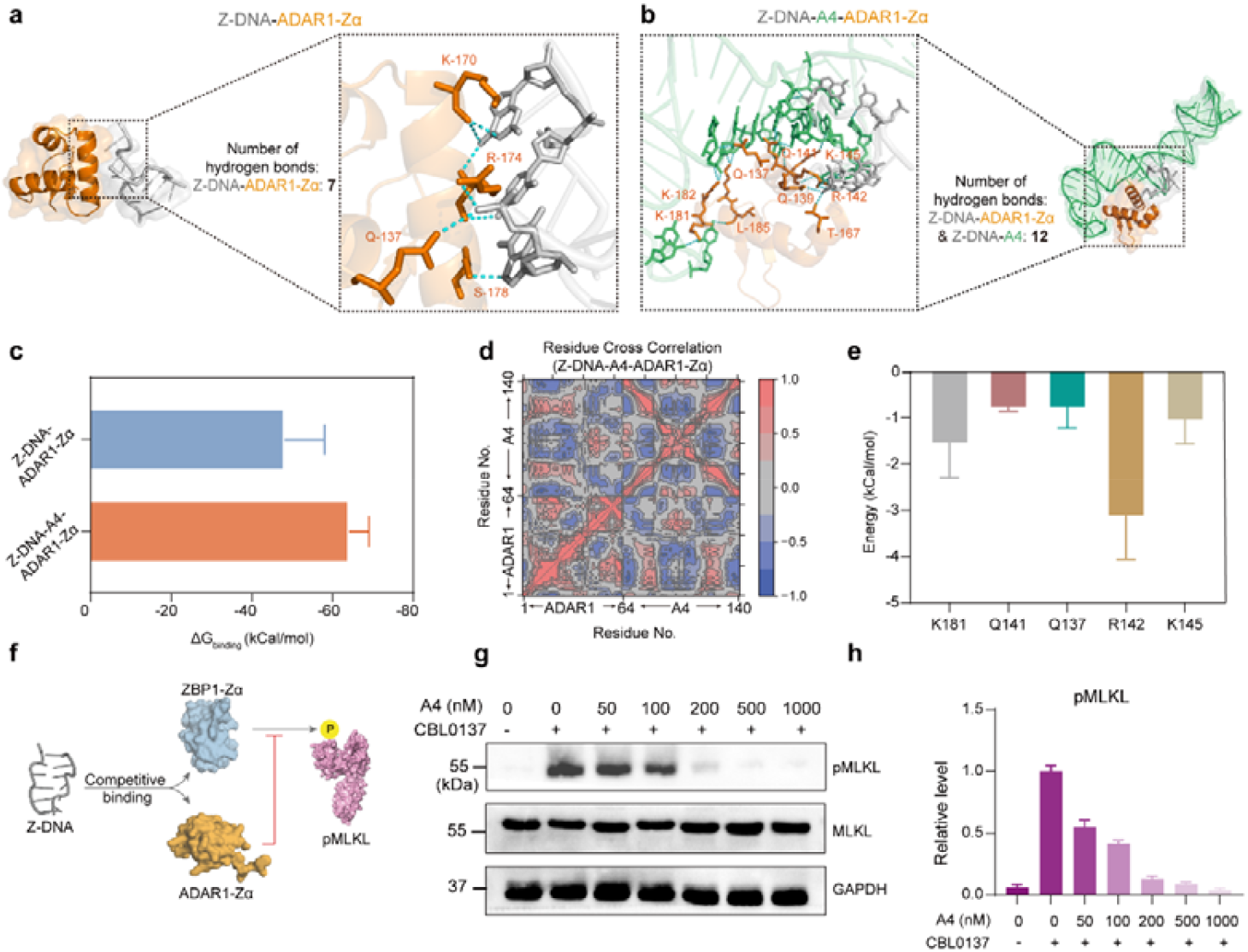
Mechanistic basis of A4-mediated enhancement of ADAR1–Z-DNA binding. **(a)** Close-up of the hydrogen bonding interactions between ADAR1-Zα and Z-DNA. **(b)** Close-up of the hydrogen bonding interactions between ADAR1-Zα, A4 and Z-DNA. (**c**) Comparison of calculated binding energies between the Z-DNA–ADAR1-Zα binary complex and the Z-DNA–A4–ADAR1-Zα ternary complex. (**d**) Residue-level cross-correlation matrix illustrating the dynamic coupling between the ADAR1-Zα domain (residues 1–64) and A4 (nucleotides 65–140) in the presence of Z-DNA. (**e**) Decomposition of binding energy contributions from key ADAR1-Zα residues (K181, Q141, Q137, R142, K145) within the ternary complex. (**f**) Schematic model of competitive Z-DNA sensing by ADAR1 and ZBP1. Z-DNA promotes ZBP1-triggered necroptosis via MLKL phosphorylation (pMLKL). Binding of ADAR1-Zα to Z-DNA competitively inhibits ZBP1 activation, thereby suppressing pMLKL-dependent necroptosis. (**g**) Western blot showing dose-dependent reduction of pMLKL levels in A549 cells treated with increasing concentrations of A4 for 24 hours. (**h**) Quantification of pMLKL expression from (g). Data are presented as mean ± s.d.; n = 3 biologically independent samples.

Further binding energy calculations demonstrated that the Z-DNA-A4-ADAR1-Zα ternary complex exhibits significantly enhanced stability compared to the binary Z-DNA-ADAR1-Zα complex, as evidenced by lower binding energies (Figure 2c). The dynamic cross-correlation matrix demonstrated strong long-range correlations between ADAR1-Zα and A4, indicating highly coupled motions in both the presence and absence of Z-DNA (Figure 2d, S5d). This concerted dynamic behavior likely reflects a capacity for coordinated conformational adjustments, which would facilitate the formation and maintenance of a stable complex. Molecular dynamics simulations of A4 in its free versus ADAR1-Zα-bound states revealed critical structural rearrangements (Figures S5e-f). While Z-DNA does not interact with the target binding region of A4 in the free state, the ADAR1-Zα-bound conformation enabled five additional hydrogen bonds with Z-DNA’s functional region, consistent with our ternary complex simulations (Figure 2b). This demonstrated that ADAR1-Zα binding induces the functional conformation required for A4’s enhancing activity. Structural modeling identified key hydrogen-bonding residues in ADAR1-Zα critical for A4 interaction (Figure 2e). Site-directed mutagenesis of these residues to alanine significantly reduced Z-DNA binding affinity in SPR experiments (Figure S5g), validating our computational predictions and confirming the mechanistic basis of A4’s activity.

Additionally, as previous studies indicated before, enhancing ADAR1 to bind with Z-DNA will competitively inhibiting ZBP1, thus attenuating necroptosis in the cell (Figure 2f).^14, 27, 28^ To validate the inhibitory function of A4 in a cellular context, we used CBL0137 to induce Z-DNA formation in A549 cells (confirmed by circular dichroism spectroscopy, Figure S5h) and transfected A4 into the cells.^14^ As shown in Figure 2g-h, Western blot analysis revealed a concentration-dependent decrease in phosphorylated MLKL (pMLKL) levels, indicating that A4 effectively attenuated CBL0137-induced necroptosis. In contrast, when the A4 Loop2+3+4 mutant was used as a negative control (NC), no significant change in pMLKL levels was observed, confirming that the inhibitory effect specifically depends on A4’s intact binding structure (Figure S5i and S5j). Further experiment also proved that A4 doesn’t affect ADAR1’s endogenous RNA-editing activity, showing its specificity in the enhancement of ADAR1’s binding with Z-DNA (Figure S6). In conclusion, both experimental and computational evidence demonstrated that A4 effectively suppresses programmed necrosis by forming a stable hydrogen-bonding network with ADAR1-Zα. This interaction enhances ADAR1-Zα’s binding affinity for Z-DNA while competitively inhibiting ZBP1 from accessing Z-DNA, thereby providing a mechanistic basis for A4’s cellular protective function.

### Mannosylation of the A4 aptamer for specific lung-targeted delivery

To further investigate the efficacy of the enhancer in suppressing excessive inflammation, we employed a pneumonia model induced by influenza virus. Immunocytes, including macrophages and dendritic cells (DC), play a crucial role in immune defense and inflammation regulation, are particularly susceptible to influenza infection.^29^ Previous studies have shown that modifying target molecules with mannose enables specific binding to the high levels of mannose receptor (CD206) on the surface of immunocytes, including alveolar macrophages and dendritic cells (DC), thereby achieving efficient and specific targeting of immunocytes in the lung.^30-33^ Therefore, we modified the enhancer A4 with mannose to selective target and inhibit ZBP1-mediated necroptosis in the pulmonary lesions of the infected model (Figure 3a and 3b). The synthesis process began with treating acetylated mannose with thiopropionic acid in the presence of boron trifluoride diethyl etherate at room temperature for 5 hours to afford compound 1a (Figure S7a)^34^. After mild deacetylation under basic conditions, the resulting mannose thio-glycoside was conjugated to the terminal amino group of A4 via an NHS/EDC coupling reaction, yielding mannosylated A4 (Man-A4) (Figure 3a). All intermediates and the final product were extensively characterized using nuclear magnetic resonance (NMR) spectroscopy and high-resolution mass spectrometry (HRMS), with detailed characterization data available in the supplementary materials (Figure S8-S10). Next, we confirmed the successful synthesis of Man-A4 using 10% polyacrylamide gel electrophoresis (PAGE) and infrared spectroscopy. The PAGE results showed a notable shift in the migration rate of A4 following mannose conjugation, while distinct changes in the infrared absorption spectrum further validated successful coupling (Figure S7b and S7c).

**Figure 3.**
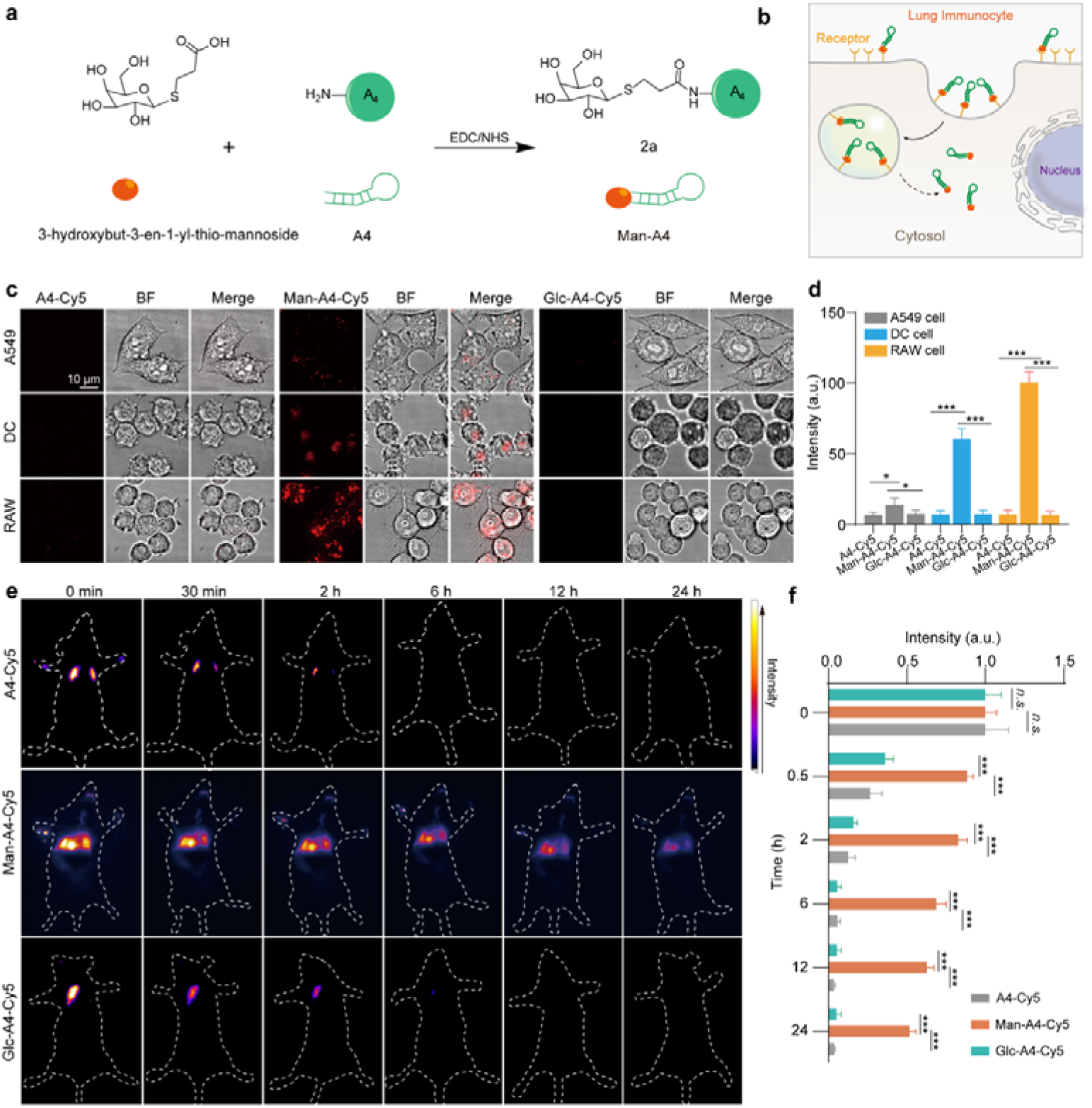
Lung-targeted delivery enabled by mannosylation of the A4 aptamer. (**a**) Reaction scheme for conjugation of A4 with 3-hydroxybut-3-en-1-yl-thio-mannoside to generate Man-A4. (**b**) Schematic model illustrating Man-A4 uptake by lung immunocytes via mannose receptor-mediated endocytosis. (**c**) Representative confocal images of A549 epithelial cells, dendritic cells, and RAW 264.7 macrophages after 6-hour incubation with Cy5-labeled A4, Man-A4, or glucosylated control (Glc-A4-Cy5). (**d**) Quantification of cellular fluorescence intensity from (c). Data are presented as mean ± s.d. (n = 10). (**e**) *In vivo* fluorescence imaging of mice at selected time points after intratracheal administration of A4-Cy5, Man-A4-Cy5, and Glc-A4-Cy5. (**f**) Time-dependent quantification of lung fluorescence intensity from (e). Data are presented as mean ± s.d. (n = 3). Statistical significance was determined by Student’s t-test (*P < 0.05, **P < 0.01, ***P < 0.001).

To evaluate the selective uptake of mannose-conjugated aptamers by immune cells, we systematically compared three Cy5-labeled compounds: unmodified A4-Cy5 to evaluate intrinsic cellular affinity; glucosylated A4-Cy5 (Glc-A4-Cy5) as a carbohydrate conjugation control lacking targeting specificity (characterized by mass spectrometry in Figure S8b); and the fully functional Man-A4-Cy5 combining both targeting and functional elements. These constructs were incubated in parallel with A549 epithelial cells, dendritic cells, and RAW 264.7 macrophages to assess cell-type-specific uptake patterns. Confocal imaging revealed a strict correlation between molecular design and internalization efficiency (Figure 3c and 3d). The control constructs (A4-Cy5 and Glc-A5-Cy5) exhibited minimal fluorescence across all cell types, indicating negligible non-specific uptake. In contrast, Man-A4-Cy5 demonstrated robust and selective internalization exclusively in immune cells, showing intense cytoplasmic fluorescence in both RAW 264.7 macrophages and dendritic cells, while remaining virtually undetectable in A549 epithelial cells. This distinct uptake pattern establishes that mannose modification is essential for immune cell targeting, as demonstrated by the failure of non-mannosylated controls to achieve significant cellular internalization. The specific and efficient delivery of Man-A4-Cy5 to immune cells provides a solid experimental basis for developing targeted immunotherapy strategies.

To systematically evaluate the pulmonary targeting and retention conferred by mannose modification, we conducted a pharmacokinetic analysis comparing the three Cy5-labeled constructs using *in vivo* fluorescence imaging (Figure 3e). Following intratracheal administration, all compounds initially accumulated in the lungs. Prolonged lung retention was observed exclusively for Man-A4-Cy5, which maintained a fluorescence intensity over 5-fold higher than the controls at 24 hours, in stark contrast to the rapid clearance (nearly 90% signal loss within 2 hours) seen with both A4-Cy5 and Glc-A4-Cy5 (Figure 3f). The persistence of a detectable signal for over 24 hours confirmed that mannose-specific modification, beyond general carbohydrate conjugation, is essential for enhanced retention, an effect potentially mediated by CD206. To further characterize the systemic disposition of the aptamer, we analyzed organ distribution at 48 hours after administration. As the pulmonary fluorescence diminished, a corresponding signal increase was observed in the liver and kidneys (Figure S7d and S7e), indicating that after initial lung engagement, Man-A4-Cy5 undergoes systemic redistribution and is ultimately cleared via hepatobiliary and renal pathways, consistent with typical elimination routes for DNA-based therapeutics.^35, 36^

### Cellular validation of the therapeutic potential of Man-A4

To confirm that Man-A4 attenuates inflammation and cell necrosis by specifically inhibiting ZBP1 and its downstream signaling pathways, we analyzed key necroptosis-associated proteins by Western blotting. The results showed that while total protein levels of RIPK3 and MLKL remained unchanged, their phosphorylation levels were significantly reduced. In contrast, the expression levels of ZBP1 and Caspase-8 were not affected (Figure 4a, S11). These findings suggested that Man-A4 suppressed necroptosis by inhibiting ZBP1 activity rather than its expression, thereby preventing the activation of RIPK3 and MLKL. Furthermore, Man-A4 did not affect Caspase-8 cleavage, which is associated with apoptosis, confirming its specific targeting of the necroptotic pathway. Dose-response analysis revealed that Man-A4 reduced pRIPK3 and pMLKL levels with half maximal effective concentration (EC_50_) values of 56.8 nM and 40.5 nM, respectively (Figure 4b). In contrast, Man-NC produced no significant response across the concentration range tested for phosphorylation levels of RIPK3 and MLKL (Figure S12). Based on this efficacy profile, 100 nM Man-A4 was chosen for subsequent in vitro experiments. Collectively, these results indicated that Man-A4 selectively inhibiting the necroptotic signaling pathway.

**Figure 4.**
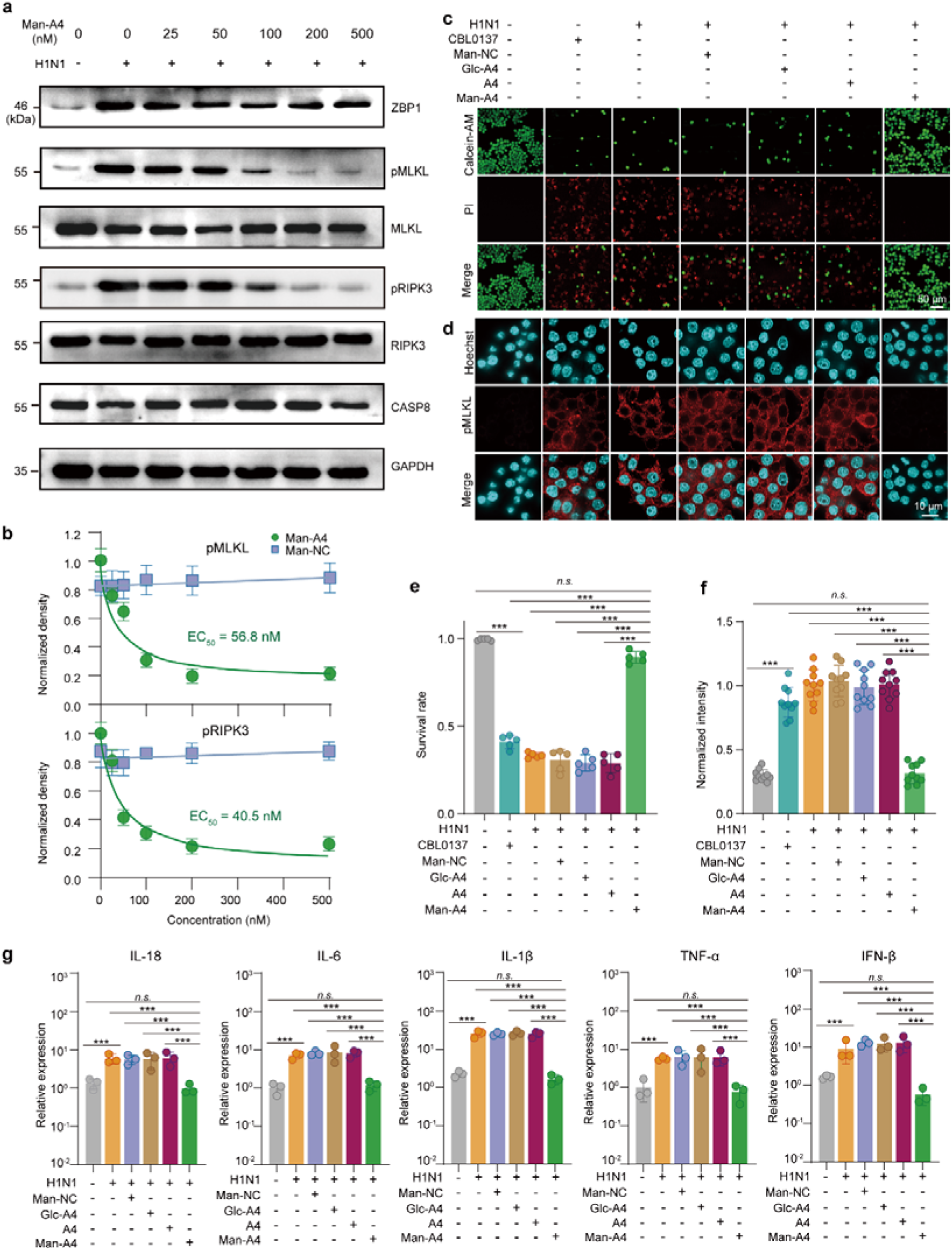
Mechanistic validation of Man-A4 as a competitive ZBP1 inhibitor suppressing necroptotic signaling. (**a**) Western blot analysis of necroptosis-related proteins (ZBP1, MLKL, pMLKL, RIPK3, pRIPK3, Caspase-8) in RAW 264.7 cells treated with increasing concentrations of Man-A4. (**b**) Dose–response curves of pMLKL and pRIPK3 inhibition by Man-A4. EC_50_ values were determined as 56.8 nM and 40.5 nM, respectively. Data are derived from (a) and Figure S11a. (**c**) Representative live/dead staining of the cells treated with Man-A4 (100 nM) and control constructs (Man-NC, Glc-A4, unmodified A4). (**d**) Immunofluorescence images showing pMLKL localization and intensity in RAW 264.7 cells across treatment groups. (**e**) Quantification of cell viability from (c) (n = 5). (**f**) Quantitative analysis of pMLKL fluorescence intensity from (d) (n = 10). (**g**) qRT-PCR analysis of inflammatory cytokine mRNA levels (IL-18, IL-1β, IL-6, TNF-α, IFN-β) in virus-infected cells under each condition (n = 3). Data are presented as mean ± s.d. Significance was assessed by Student’s *t*-test: P < 0.05, *P < 0.01, **P < 0.001.

To systematically evaluate the therapeutic potential of Man-A4 against virus-induced cellular damage, we established two independent inflammation models using CBL0137 treatment and viral infection, and performed comprehensive analysis through live/dead cell staining and immunofluorescence assays. Both models recapitulated key features of virus-induced cytotoxicity, with cell viability declining dramatically to 41.1% in the CBL0137 model and 33.4% in the viral infection model (Figure 4c and 4e). These results confirmed the severe cellular damage induced by both stimuli. Strikingly, Man-A4 treatment demonstrated remarkable protective efficacy, restoring cell viability to 89.5% despite the challenging inflammatory environment. Importantly, this protective effect was specific to the functional Man-A4 construct, as control variants including Man-NC, Glc-A4, and unmodified A4 showed minimal protective capacity under identical conditions.

To elucidate the molecular mechanism underlying Man-A4-mediated cytoprotection, we performed immunofluorescence analysis of phosphorylated MLKL (pMLKL), the core executor of necroptosis, to evaluate its subcellular localization and expression dynamics. The untreated control cells exhibited low baseline pMLKL signals, reflecting minimal necroptotic activation under physiological conditions. In contrast, both CBL0137 treatment and viral infection markedly enhanced pMLKL intensity in the cytosol, demonstrating robust pathway activation. Notably, Man-A4 treatment significantly suppressed this increase, reducing pMLKL to levels comparable to the untreated cells, whereas the Man-NC, Glc-A4, and unmodified A4 controls showed no significant suppression of pMLKL signal (Figures 4d and 4f). The specific inhibitory effect observed only with Man-A4 underscores its unique activity in regulating necroptosis. Together, these results demonstrated that Man-A4 attenuates virus-induced necroptosis by selectively inhibiting MLKL phosphorylation, thereby maintaining cellular viability.

Further qRT-PCR analysis confirmed that viral infection significantly upregulated multiple inflammatory mediators, including IL-18, IL-6, IL-1β, TNF-α, and IFN-β (Figure 4g), indicative of a strong inflammatory reaction. TNF-α functioned as a primary initiator of the inflammatory cascade, while IFN-β served as a key mediator of antiviral defense. The rise in IL-18 reflected its involvement in pro-inflammatory signaling, while marked increases in IL-6 and IL-1β underscored the severity of the inflammatory response. Treatment with Man-A4 substantially reduced the expression of these cytokines, in contrast to control constructs (Man-NC, Glc-A4, and unmodified A4), which showed no significant suppressive effect. These findings supported the conclusion that Man-A4 inhibits the necroptotic signaling pathway, leading to reduced production of pro-inflammatory cytokines. The data further suggested that Man-A4 may play a dual role in modulating inflammation—through suppression of cytokine expression and possible restoration of immune homeostasis via additional signaling pathways. In summary, Man-A4 effectively alleviated virus-triggered inflammation and represents a promising therapeutic candidate for the management of inflammatory responses in viral infections.

### *In vivo* evaluation of the anti-inflammatory effects of Man-A4

To comprehensively evaluate the anti-inflammatory efficacy and therapeutic potential of Man-A4 in influenza A virus (IAV) infection, we established a lethal murine model by intranasally challenging Balb/c mice with 10 × TCID□□ of IAV. Beginning 24 hours post-infection, we administered Man-A4 systemically once daily for a total of 4 days. Through this rigorous experimental design, we systematically investigated the dose-response relationship of Man-A4 across a concentration range of 5-30 mg/kg and meticulously monitored survival rates (Figure 5a). Our results revealed a well-defined dose-dependent protection pattern. At the lowest tested dose of 5 mg/kg, Man-A4 demonstrated significant therapeutic efficacy by rescuing 30% of infected mice from mortality. The protective effect showed a clear positive correlation with dosage increases, reaching its optimal efficacy at 20 mg/kg. However, when we further escalated the dosage to 30 mg/kg, no additional survival benefit was observed, suggesting that 20 mg/kg represents the saturation point for Man-A4’s therapeutic effect in this model (Figure 5b). In a parallel experiment designed to better mimic clinical scenarios, we employed a less lethal but clinically more relevant H1N1 challenge (5 × TCID□□). Remarkably, in this model, Man-A4 at an intermediate dose of 10 mg/kg achieved complete (100%) protection, preventing all mortality events (Figure 5c). To confirm the specificity of Man-A4’s therapeutic action, we conducted extensive control experiments. Treatments with 30 mg/kg of Man-NC, Glc-A4, or unmodified A4 all showed survival rates statistically indistinguishable from the PBS control group. This comprehensive control validation firmly established that the observed protection specifically requires the integrated functional mannosylated aptamer structure, highlighting the unique therapeutic profile of Man-A4. The clear dose-response relationship, combined with the complete protection observed at lower viral loads and the stringent specificity controls, provides compelling evidence for Man-A4’s potential as a targeted therapeutic agent against virus-induced inflammation and mortality.

**Figure 5.**
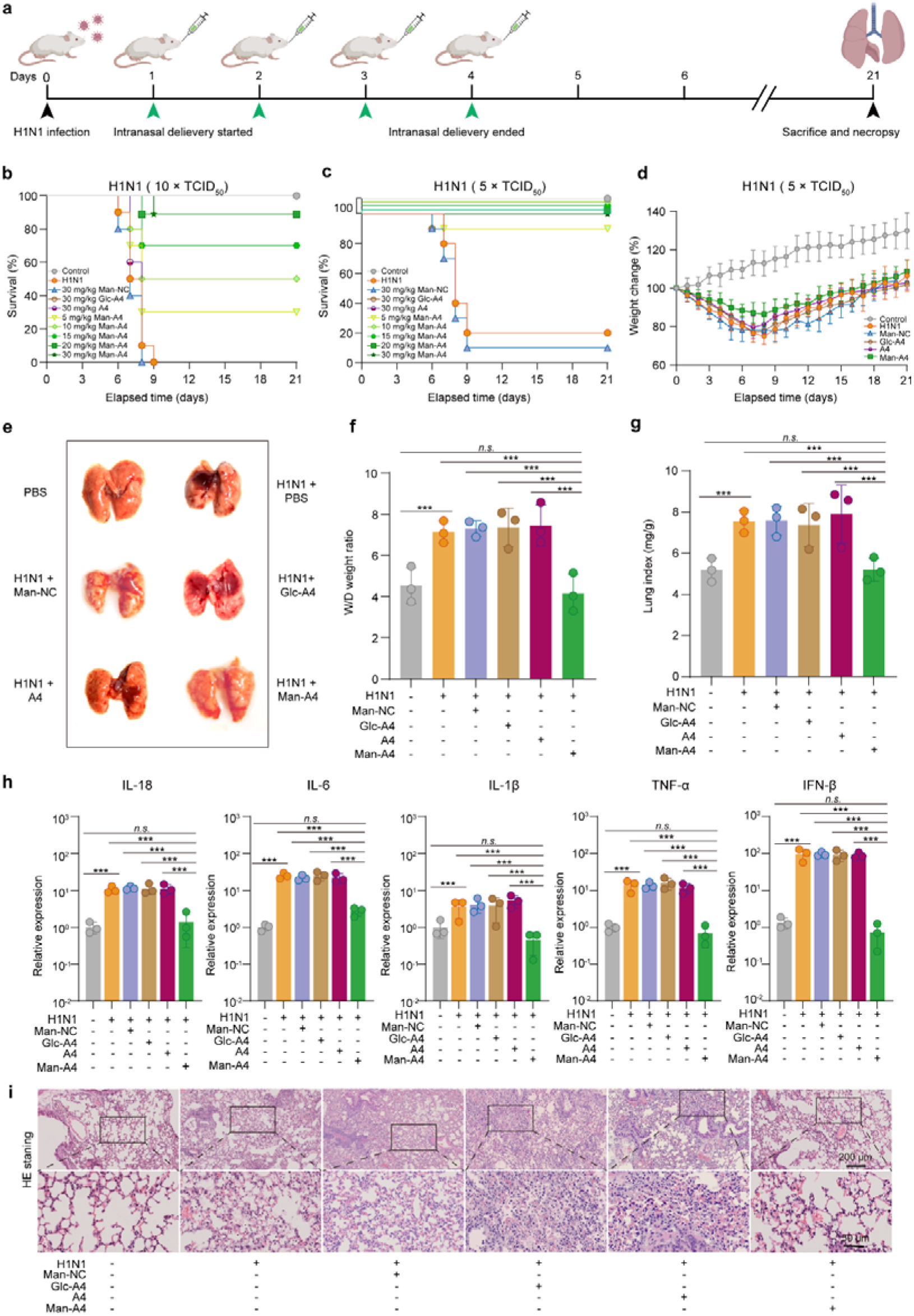
*In vivo* evaluation of the anti-inflammatory efficacy of Man-A4 in IAV-infected mice. (**a**) Schematic diagram of the experimental timeline for viral infection and treatment. (**b**) Survival curves of mice infected with a lethal dose of H1N1 (10 × TCID□□) and treated intratracheally once daily with PBS, Man-NC, Glc-A4, unmodified A4 (30 mg/kg), or Man-A4 at the indicated doses (5–30 mg/kg; n = 20 per group). (**c**) Survival analysis following infection with a moderate viral inoculum (5 × TCID□□) and treatment as in (b). (**d**) Body weight changes in H1N1-infected mice treated with PBS or Man-A4 (20 mg/kg) once daily (n = 20). (**e**) Representative lung images from each group showing the extent of inflammation. (**f**) Lung wet-to-dry weight ratios across experimental groups (n = 3). (**g**) Lung index values reflecting pulmonary edema and inflammation (n = 3). (**h**) qRT-PCR analysis of inflammatory cytokine mRNA levels in lung tissues. (**i**) Hematoxylin and eosin (H&E) stained lung sections with magnified views of regions outlined in black, illustrating inflammatory cell infiltration and alveolar integrity. Data are presented as mean ± s.d. Significance was determined by Student’s t-test: *P < 0.05, **P < 0.01, **P < 0.001; n.s., not significant. Scale bars are shown in the respective panels.

Man-A4 treatment demonstrated significant protective effects in IAV-infected mice, not only delaying the onset of weight loss but also substantially reducing the overall magnitude of weight reduction compared to vehicle-treated controls (Figure 5d). Gross anatomical examination of lung tissues revealed severe pulmonary edema and congestion in virus-infected animals, whereas no apparent inflammation was observed in uninfected controls. Importantly, Man-A4 treatment markedly attenuated lung inflammation, with tissue appearance approaching normal morphology. In contrast, lungs from Man-NC, Glc-A4, and unmodified A4 treatment groups exhibited pronounced inflammatory responses comparable to the virus-infected group (Figure 5e). To quantitatively assess pulmonary edema, we measured the lung wet-to-dry weight ratio (Figure 5f). Both H1N1-infected mice and those treated with control constructs (Man-NC, Glc-A4, and A4) showed significantly elevated ratios, indicating severe lung inflammation and edema formation. Conversely, Man-A4 treatment effectively normalized this parameter. These findings were further corroborated by lung index measurements, which demonstrated significant increases in all control groups but returned to normal range following Man-A4 administration (Figure 5g). At the molecular level, qRT-PCR analysis revealed that viral infection markedly upregulated pulmonary expression of pro-inflammatory cytokines (IL-18, IL-6, IL-1β, TNF-α, and INF-β) in the lung tissues of both infected mice and groups receiving inactive controls. Man-A4 treatment significantly suppressed this cytokine induction to nearly normal level (Figure 5h). Consistent with these observations, western blot analysis confirmed that pMLKL levels in Man-A4-treated lungs were substantially lower than those in PBS, Man-NC, Glc-A4, and A4 treatment groups (Figure S13a and S13b). Collectively, these results indicated that Man-A4 effectively inhibits pro-inflammatory cytokine production and modulates immune responses through activation of ADAR1 and subsequent suppression of the ZBP1-mediated necroptotic signaling pathway.

Histopathological examination through hematoxylin-eosin (H&E) staining further validated the therapeutic efficacy of Man-A4 against viral pneumonia (Figure 5i and S13c)). Lung tissues from the viral infection group and control groups (Man-NC, Glc-A4, and A4) displayed extensive inflammatory cell infiltration and severe alveolar structural damage. In contrast, Man-A4 treatment significantly ameliorated these pathological alterations, with substantially diminished inflammatory cell accumulation and restored tissue architecture. Terminal deoxynucleotidyl transferase dUTP nick end labeling (TUNEL) staining provided further mechanistic insights, revealing extensive apoptotic cell death in lung sections from the virus-infected group and control construct-treated groups. Notably, Man-A4 treatment dramatically reduced TUNEL-positive signals to levels comparable with uninfected controls (Figure S13d and S13e), demonstrating its potent anti-apoptotic activity *in vivo*. Safety assessment across major organs revealed no Man-A4-induced pathological changes in cardiac, hepatic, splenic, or renal tissues (Figure S13f), supporting its favorable safety profile. In summary, our *in vivo* findings robustly demonstrated that Man-A4 effectively activates ADAR1 while inhibiting ZBP1-mediated signaling, resulting in comprehensive therapeutic outcomes against viral pneumonia.

## Conclusion

This study successfully developed a SELEX-HTCFQ screening platform that incorporated competitive fluorescence quenching to enable efficient identification of functional domain-specific aptamers. Using this platform, we identified the bifunctional aptamer A4, which specifically enhanced ADAR1-Zα binding to Z-DNA while competitively inhibiting ZBP1-Zα interaction with Z-DNA. This dual mechanism effectively suppressed pathological inflammation in both cellular and animal models without compromising fundamental immune functions. Structural characterization revealed that A4 formed an extensive hydrogen-bond network with ADAR1-Zα, significantly increasing its binding affinity for Z-DNA and providing a molecular basis for the observed anti-inflammatory effects. To enhance therapeutic potential, we engineered a mannosylated derivative, Man-A4, which exhibited improved pulmonary targeting and prolonged tissue retention. In viral pneumonia models, Man-A4 demonstrated potent anti-inflammatory and anti-necroptotic activity through multi-mechanistic regulation of the ADAR1-ZBP1 signaling axis. These findings not only addressed limitations of conventional SELEX approaches but also established ADAR1 enhancers as a novel therapeutic strategy for precise pathway modulation.

Compared to the SELEX methods that primarily targeted extracellular epitopes,^24, 35, 37^ our modular conjugation strategy introduced a paradigm shift by successfully decoupling the target recognition function from cellular delivery mechanisms. This strategic separation enabled the engineering of aptamers with enhanced therapeutic capabilities, significantly expanding their functional scope from conventional diagnostic applications to sophisticated intracellular therapeutics. The mannosylation approach exemplified this advancement, bestowing cell-penetrating properties while preserving the intrinsic molecular recognition characteristics of the aptamer. Despite these promising developments, several formidable challenges persisted in the clinical translation of aptamer-based therapeutics. The inherent susceptibility to nuclease degradation posed a significant barrier to therapeutic stability, while rapid renal clearance mechanisms limited systemic exposure and required frequent dosing regimens. Furthermore, achieving efficient tissue-specific delivery remained an ongoing challenge that demanded innovative solutions. These limitations highlighted the necessity for continued investigation into advanced conjugation chemistries, novel delivery platforms, and optimized formulation strategies to enhance the pharmacokinetic profiles and therapeutic indices of aptamer-based medicines.

Collectively, our findings represented a significant departure from conventional therapeutic approaches that relied on simplistic inhibition or activation of biological targets. Instead, we demonstrated the feasibility of context-dependent pathway fine-tuning through precise biomolecular regulation. This approach maintained the delicate balance of physiological homeostasis while effectively countering disease processes, particularly in complex inflammatory conditions. The study established a robust foundation for the continued development of aptamer-based precision medicines, providing both methodological insights and conceptual advances. By demonstrating the therapeutic potential of ADAR1 enhancers and establishing a generalizable platform for intracellular aptamer delivery, this work opened new avenues for treating a broad spectrum of diseases that require precise modulation of cellular pathways rather than complete pathway blockade or activation.

## Supporting Information

Additional experimental details, materials, methods, animals used in this study, experimental section, Figures S1−S13, Table S1-S4, DNA sequences used, and references.

## Acknowledgements

This work was supported by the National Key Research and Development Program of China (Nos. 2024YFA1210003), and Tianjin Natural Science Foundation (Nos. 24JCZDJC01240, 23JCZDJC01240, and 23JCYBJC01880).

## Conflict of Interest

The authors declare no conflict of interest.

## Notes

### Competing Interest Statement

The authors have declared no competing interest.

### Summary of Updates

Figure 3, a typo on the chemical reaction was corrected; all the "INF-β" was revised to "IFN-β" in Figure 4, 5, and the text.

