## supplemental material for "SELEX-HTCFQ Platform: Developing DNA Enhancers of ADAR1 to Suppress ZBP1-Dependent Immunopathology"

### Materials and Methods

#### Materials

Dulbecco's modified Eagle's medium (DMEM) was obtained from Gibco (Grand Island, NY, USA). The One-step TUNEL Apoptosis Assay Kit with green fluorescence detection was supplied by Abbkine Scientific Co., Ltd. Polyclonal antibodies targeting ADAR1, MLKL, pMLKL, RIPK3, pRIPK3, Caspase-8, and GAPDH were acquired from Signalway Antibody LLC (USA). All DNA sequences were custom-synthesized by Tsingke Biotechnology Co., Ltd. (Beijing, China). The small molecule compound CBL0137 (Catalog No. C910450) was procured from MACKLIN.

#### Protein Expression and Purification

The plasmid encoding the His-tagged ADAR1-Zα domain was transformed into E. coli BL21(AI) cells. A single colony was selected and cultured overnight at 37 °C in LB medium supplemented with the appropriate antibiotic. The following day, the overnight culture was diluted 1:100 into fresh LB medium and grown at 37 °C until the OD₆₀₀ reached 0.6–0.8. Protein expression was induced by adding 0.1% L‑arabinose and 1 mM IPTG, followed by incubation at 20 °C for 8–12 h. Cells were harvested by centrifugation at 4 °C and resuspended in lysis buffer (50 mM Tris‑HCl, 300 mM NaCl, 10% glycerol, pH 8.0). After resuspension, the cells were lysed by sonication on ice, and the lysate was clarified by centrifugation. The soluble supernatant was loaded onto a Ni‑NTA resin column pre‑equilibrated with binding buffer. The column was washed with wash buffer (50 mM Tris‑HCl, 300 mM NaCl, 20 mM imidazole, pH 8.0) to remove nonspecifically bound proteins. Target protein was eluted with elution buffer containing 500 mM imidazole. The purified protein was quantified using a BCA protein assay kit, and its purity and yield were evaluated by SDS‑PAGE.

#### SELEX Technology for Aptamer Selection

The SELEX procedure for aptamer selection comprised multiple critical stages involving primary library screening, secondary library preparation, and iterative rounds of positive and negative selection to isolate high-affinity aptamers.^1^ The process was initiated by selecting a single-stranded DNA (ssDNA) library and preparing magnetic beads. The complete sequence of the library was: 5'-TTCAGCACTCCACGCATAGCNNNNNNNNNNNNNNNNNNNNNNNNNNNNNNNNNNNNCCTATGCGTGCTACCGTGAA -3', where "N" represents an equimolar mixture of A, T, G, and C. The central random region consisted of 36 nucleotides (nt). The fixed 5' and 3' primer binding sites were 20 nt and 20 nt in length, respectively. Specifically, 10 μL of tosyl-activated magnetic beads (TMB) were mixed with 50 μL of 0.1 mg/mL ADAR1‑Zα protein solution in a suitable buffer and incubated overnight at 4 °C to immobilize the target protein. After incubation, the beads were washed once with PBS and twice with PBST to remove unbound protein. The TMB‑protein complexes were then incubated with 1 OD (100 μmol/L) of the ssDNA library for 2 h at 4 °C to enable binding. Subsequently, bound ssDNA molecules were recovered through sequential washing and thermal elution, followed by PCR amplification. The amplified products were verified by 4% agarose gel electrophoresis to confirm amplification specificity.

The PCR products were purified using a column-based PCR purification kit, and the concentration of the resulting double-stranded DNA (dsDNA) was measured with a microplate reader. The purified dsDNA was incubated with streptavidin magnetic beads at a ratio of 100 ng dsDNA per 5 μL beads for 30 minutes. To generate single-stranded DNA, the dsDNA was denatured by adding NaOH, and the unbound ssDNA was subsequently eluted. The eluate was neutralized with NaH₂PO₄ to obtain the secondary ssDNA library. For subsequent selection rounds, this secondary ssDNA library served as the input for both positive and negative selection procedures. In positive selection, the library was incubated with the immobilized target protein, followed by washing steps to remove unbound sequences. Negative selection was performed using beads without conjugated target protein to eliminate nonspecific binders. Through ten iterative rounds of positive and negative selection, aptamers with high specificity and affinity for the target protein were effectively enriched. This iterative screening process progressively selected ssDNA aptamers with strong binding capabilities, ultimately yielding candidate aptamers for downstream characterization.

#### HTCFQ Assay for Screening ADAR1 Enhancers

We employed a fluorescence quenching-based methodology to investigate protein-Z-DNA interactions and aptamer-mediated regulatory effects. The experimental setup involved a 3'-BHQ2-labeled DNA strand that was converted into Z-DNA-BHQ2 probes through incubation with 10 μM CBL0137 for 30 minutes at room temperature. The fluorescent fusion constructs ADAR1-Zα-mRFP1 and ZBP1-Zα-mOrange were prepared using established expression and purification protocols. The initial fluorescence intensities (designated as F_O_ for ZBP1-Zα-mOrange and F_R_ for ADAR1-Zα-mRFP1) were measured at 562 nm and 607 nm, respectively, after incubating the two proteins (100 nM each) together for 5 minutes at room temperature. To determine the optimal working concentration of Z-DNA-BHQ2, ADAR1-Zα-mRFP1 or ZBP1-Zα-mOrange (100 nM) was incubated with varying concentrations of Z-DNA-BHQ2 for 5 minutes at room temperature. The fluorescence intensities in the absence of aptamer (F_O-blank_ and F_R-blank_) were measured at 562 nm and 607 nm, respectively. The BHQ2-induced fluorescence quenching was calculated as ΔF_O-blank_ = F_O_ - F_O-blank_ and ΔF_R-blank_ = F_R_ - F_R-blank_. Plots of ΔF_o-blank_ and ΔF_R-blank_ versus Z-DNA-BHQ2 concentration were generated, and the concentration corresponding to approximately half-maximal quenching (15 nM in this study) was selected for subsequent experiments.

For binding measurements, ADAR1-Zα-mRFP1 or ZBP1-Zα-mOrange (100 nM) was incubated with individual aptamers (100 nM) and 15 nM Z-DNA-BHQ2 for 5 minutes at room temperature. Fluorescence intensities were measured at 562 nm and 607 nm, and the apparent fluorescence changes (ΔF_O_ and ΔF_R_) were calculated by comparison with the initial F_O_ and F_R_ values. These apparent signals comprise three components: specific BHQ2-mediated quenching (ΔF_mOrange_ or ΔF_mRFP1_), nonspecific quenching (ΔF_n_), and spectral bleed-through between fluorescence channels (ΔF_b_). To isolate the specific quenching signal resulting from protein-BHQ2 binding, control experiments were performed to determine both correction factors. Nonspecific quenching (ΔF_n_), caused by transient proximity between fluorophores and the quencher, was quantified by incubating 100 nM mRFP1 or mOrange with 100 nM aptamer A4 and 15 nM Z-DNA-BHQ2. The resulting fluorescence changes at 562 nm and 607 nm yielded ΔF_nO_ and ΔF_nR_, respectively. Spectral bleed-through (ΔF_b_), arising from detection of one fluorophore's signal at the other's emission wavelength, was assessed by measuring the fluorescence change at 562 nm when 100 nM ADAR1-Zα-mRFP1 and 100 nM aptamer A4 were incubated with 15 nM Z-DNA-BHQ2 (ΔF_bO_), and similarly at 607 nm using ZBP1-Zα-mOrange to obtain ΔF_bR_. The corrected specific fluorescence changes were then calculated using the equations ΔF_mOrange_ = ΔF_O_ - ΔF_nO_- ΔF_bO_ and ΔF_mRFP1_ = ΔF_R_ - ΔF_nR_ - ΔF_bR_. The ΔF_mRFP1_/ΔF_mOrange_ ratio was computed for all aptamers, and the aptamer with the highest ratio was selected for further investigation.

The dissociation constants (K_D_) for Z-DNA binding under different conditions were determined through systematic titration experiments. For measuring the K_D_ of Z-DNA binding to individual proteins, ADAR1-Zα-mRFP1 or ZBP1-Zα-mOrange (100 nM) was incubated with increasing concentrations of Z-DNA-BHQ2 in a 96-well plate for 5 minutes at room temperature. Fluorescence intensity changes (ΔF) were monitored at their respective detection wavelengths (607 nm for mRFP1 and 562 nm for mOrange). Since this single-protein system did not involve mixed fluorescent proteins, spectral bleed-through correction was unnecessary, and only nonspecific quenching required correction. The nonspecific quenching components (ΔF_nO_ and ΔF_nR_) were determined by parallel experiments using 100 nM mRFP1 or mOrange with 100 nM A4 and the same Z-DNA-BHQ2 concentration series. The corrected specific signals were calculated as ΔF_mOrange_ = ΔF_O_ - ΔF_nO_ and ΔF_mRFP1_ = ΔF_R_ - ΔF_nR_. The maximum values (ΔF_max_) from ΔF_mOrange_ and ΔF_mRFP1_ datasets were used to normalize the data, and binding curves were generated by plotting ΔF/ΔF_max_ against Z-DNA-BHQ2 concentration for curve fitting.

For determining the *K*_D_ of Z-DNA binding to protein-A4 complexes, the same methodology was applied except that ADAR1-Zα-mRFP1 (100 nM) was pre-incubated with A4 (100 nM) before the addition of Z-DNA-BHQ2. In the case of competitive *K*_D_ measurements involving mixed proteins with A4 complexes, the experimental setup required inclusion of both fluorescent proteins along with A4, necessitating the comprehensive correction formula that accounted for both nonspecific quenching and spectral bleed-through components to obtain accurate ΔF_mOrange_ and ΔF_mRFP1_ values for reliable *K*_D_ determination.

#### SPR Analysis of Aptamer-ADAR1-Zα Binding

Surface plasmon resonance (SPR) analysis was conducted using a CM5 sensor chip to characterize the binding interactions between ADAR1-Zα and various ligands. The chip surface was activated with EDC/NHS to enable covalent immobilization of ADAR1-Zα. Initial screening of 16 candidate aptamers identified A4 as the highest-affinity binder through real-time response monitoring and sensorgram analysis. For quantitative affinity determination, serial concentrations of aptamer A4 (0.01-1000 nM) and Z-DNA were injected over the functionalized surface. Equilibrium dissociation constants (*K*_D_) for both ligands binding to ADAR1-Zα were derived through global fitting of the binding curves. To further investigate the ternary complex formation, the A4-ADAR1-Zα complex was first assembled in situ on the chip surface, followed by injection of increasing Z-DNA concentrations. The *K*_D_ for Z-DNA binding to the pre-formed complex was subsequently calculated, revealing enhanced binding affinity compared to the binary interaction.

#### Molecular Dynamics Simulations

The initial structures of the ADAR1-Zα-aptamer complex and the Z-DNA-A4-ADAR1-Zα ternary complex were generated using the HDOCK server, with the highest-scoring structure selected for subsequent molecular dynamics (MD) simulations.^2^ For the ADAR1-Z-DNA complex, the structure was retrieved from the Protein Data Bank (PDB ID: 1QBJ). The starting structure for the protein-aptamer complex was also derived from the HDOCK server. The input files for the MD simulations were prepared using CHARMM-GUI.^3^

To set up the system for simulations, Na^+^ and Cl^-^ ions were added to neutralize the system and achieve an ionic strength of 0.15 M. The protein was modeled using the AMBER14SB force field, while the AMBER OL15 force field was applied to the DNA aptamer.^4, 5^ The entire system was solvated in a cubic box, ensuring a minimum distance of 1.5 nm between the solute and the box edges, with TIP3P water molecules used for hydration. The system then underwent energy minimization via the steepest descent method to remove any unfavorable contacts. Following minimization, the temperature was gradually increased from 0 K to 310.15 K using the Nose-Hoover method, followed by NVT equilibration at 310.15 K to stabilize the system.^6^ After equilibration, 200 ns of MD simulations were conducted using the NPT ensemble for each system. The temperature was maintained at 310.15 K using the V-rescale coupling method, and pressure was controlled at 1 bar with the Parrinello-Rahman coupling method.^7^ Nonbonded interactions were computed with a 9 Å cutoff, and electrostatic interactions were handled using the particle mesh Ewald (PME) algorithm.^8^ Hydrogen bonds were constrained using the LINCS algorithm.^9^ The time step was set to 2 fs, and trajectory data was collected every 10 ps for further analysis.

#### Binding Free Energy Calculation

In this study, the Molecular Mechanics/Poisson Boltzmann Surface Area (MM/PBSA) model was employed to calculate the binding free energy. Before performing the calculations with gmx_MMPBSA, the MD output trajectory was properly aligned, and periodic boundary conditions (PBC) were removed. The binding free energy (∆G_bind) was calculated using the following equation:

Δ*Gbind* = Δ*H* – *T*Δ*S*

Δ*H* = Δ*EMM* + Δ*Gsol*

Δ*EMM* = Δ*Ebonded* + Δ*Enonbonded*= (Δ*Ebond* + Δ*Eangle* + Δ*Edihedral*) + (Δ*Eele* + Δ*Evdw*)

Δ*Gsol* = Δ*Gpolar* + Δ*Gnon-polar* = Δ*GPB* + Δ*Gnon-polar*

In these equations, **∆***EMM* represents the changes in molecular mechanical energy in the gas phase. It is divided into **∆***Ebonded*, or internal energy, and **∆***Enonbonded*, which includes the van der Waals (vdW) and electrostatic (ele) interactions.

The solvation energy, **∆***Gsol*, is computed by considering both polar and nonpolar components. For the polar component, the Poisson-Boltzmann (PB) model is used, while the nonpolar component is generally approximated as proportional to the molecule’s solvent accessible surface area (SASA). The proportionality constant for the nonpolar contribution is derived from experimental solvation energies of small nonpolar molecules.^10^ In contrast, the 3D-RISM model computes both polar and nonpolar solvation components.

#### Cell Culture, Virus Propagation, and Animal Model Establishment

Human alveolar basal epithelial cells (A549), mouse bone marrow-derived dendritic cells (BMDCs), and RAW 264.7 murine macrophages (ATCC TIB-71) were maintained in Dulbecco's Modified Eagle Medium (DMEM) supplemented with 10% fetal bovine serum (FBS) and 1% penicillin-streptomycin (100 U/mL penicillin and 100 μg/mL streptomycin). All cell lines were cultured at 37 °C in a humidified 5% CO₂ atmosphere.

Influenza A virus (IAV) strain A/Puerto Rico/8/1934 (H1N1, PR8) was propagated in specific pathogen-free 10-day-old embryonated chicken eggs and purified according to established protocols^11^.

Four-week-old female Balb/c mice were sourced from Beijing Vital River Laboratory Animal Technology Co., Ltd. (Beijing, China). All animal experiments were conducted in compliance with the Chinese Regulations for the Administration of Affairs Concerning Experimental Animals and approved by the Animal Ethics Committee of Nankai University (Protocol SYXK (Jin) 2019-0003). To establish the viral pneumonia model, mice were intranasally inoculated with 10× TCID₅₀ or 5× TCID₅₀ of H1N1 virus in approximately 20 μL volume. Following viral challenge, animals were housed under standard laboratory conditions for 24 hours before subsequent experimental procedures.

#### Western Blotting

Treated cells were lysed using cell lysis buffer (Solarbio, Cat. No. BC3710). Protein concentrations were quantified with the Bradford assay (Solarbio, Cat. No. PC0010), and equal amounts of protein were separated by sodium dodecyl sulfate–polyacrylamide gel electrophoresis (SDS-PAGE). Subsequently, proteins were transferred to a polyvinylidene difluoride (PVDF) membrane (Immobilon®-PSQ, Cat. No. ISEQ00010). The membrane was blocked with 5% (w/v) bovine serum albumin (BSA; Solarbio, Cat. No. A8010) for 45 min, followed by incubation with primary antibodies (1:2000 dilution) overnight at 4°C. After washing to remove unbound antibodies, the membrane was incubated with horseradish peroxidase (HRP)-conjugated secondary antibodies (1:2000 dilution) for 1 h at room temperature. Protein bands were visualized using a Bio-Rad ChemiDoc™ MP imaging system.

#### Synthesis of Mannosylated A4 (Man-A4)

**Synthesis of 1a:** The procedure was mainly following the previously published method (Figure S5a).^12^ A solution of acetylated mannose (mannose pentaacetate, 12.82 mM) and thiopropionic acid (51.28 mM) was treated with boron trifluoride diethyl etherate (BF₃OEt₂, 19.23 mM). The reaction mixture was allowed to stand at room temperature for 5 hours to yield the thioglycoside of mannose. ^1^H NMR (400 MHz, DMSO-d6) δ 12.26 (s, 1H), 5.50 (s, 1H), 5.22–5.15 (m, 1H), 5.11 (t, J = 10.0 Hz, 1H), 5.02 (dd, J = 10.1, 3.4 Hz, 1H), 4.32–4.01 (m, 3H), 2.81 (t, J = 7.0 Hz, 2H), 2.33 (dd, J = 15.5, 7.4 Hz, 2H), 2.02 (dd, J = 36.9, 32.5 Hz, 13H). ^13^C NMR (101 MHz, DMSO-d6) δ 172.83, 170.06, 169.78–169.45, 81.69, 70.05, 69.03, 68.62, 65.65, 61.98, 34.26, 26.02, 20.48, 19.34. MS (MALDI-TOF, m/z): C_17_H_24_O_11_S^+^ [H]^+^ calcd for 436.1039, found 435.0959.

**Synthesis of 1b:** A solution of 1a (20.86 mM) in DCM and methanol (1:1) was treated with 1M sodium methoxide in methanol and stirred for about 15 min at room temperature to get a white precipitate. The mixture was concentrated in rotary vacuum evaporator followed by dissolving in aqueous methanol (20%). The solution was stirred overnight at room temperature and neutralized to pH 6 using 1M HCl. ^1^H NMR (400 MHz, MeOD) δ 5.14 (s, 1H), 3.71–3.56 (m, 4H), 3.46–3.38 (m, 2H), 2.87–2.62 (m, 6H), 2.46 (d, J = 6.4 Hz, 4H). ^13^C NMR (101 MHz, DMSO-d6) δ 174.20, 85.32, 74.78, 72.06, 67.65, 61.40, 35.90, 26.50. MS (MALDI-TOF, m/z): C_9_H_16_O_7_S^+^ [H]^+^ calcd for 268.0617, found 267.0545.

**Synthesis of 2a:** A solution of compound 1b (2.67 mg, 10 μmol) in PBS-DMSO (1:1, 500 μL) was prepared. The amine-modified aptamer (230 μg, 10 nmol) was dissolved in PBS-DMSO (1:1, 500 μL). EDC (1.18 mg, 10 μmol) was added to the solution of 1b to activate the carboxyl group. The aptamer solution was then carefully mixed with the 1b and EDC solution. The mixture was stirred at room temperature for 2-4 hours to facilitate the conjugation. After completion, the product was purified using a NAP-5 desalting column, eluting with water to afford the 2a. The successful conjugation was characterized using PAGE and IR spectroscopy.

#### Synthesis of Glucosylated A4 (Glc-A4)

D-(+)-Glucuronic acid (1.94 mg, 10 μmol) was dissolved in PBS (500 μL), and its carboxyl group was activated by the addition of EDC (1.18 mg, 10 μmol). The reaction mixture was stirred at room temperature for 30 minutes. Separately, the amine-modified aptamer A4 (230 μg, 10 nmol) was dissolved in PBS (500 μL). The activated solution was then combined with the aptamer solution, and the mixture was stirred at room temperature for 6 hours to facilitate amide bond formation. The crude product was purified using a NAP-5 desalting column with water as the eluent to yield Glc–A4.

#### Cell Uptake

A549, DC, and RAW 264.7 cells were pre-cultured and then incubated for 6 hours with one of three treatments: A4-Cy5, Glc-A4, or mannose-A4-Cy5. Following incubation, the cells were washed three times with PBS to remove unbound compounds. The intracellular distribution of the aptamers was subsequently analyzed by confocal microscopy. Imaging was performed using a 640 nm laser for excitation and a 685/40 nm band-pass filter for emission capture.

#### Immunofluorescence Staining

Following experimental treatments, cells were washed three times with PBS (3 min per wash) and fixed with 4% paraformaldehyde for 30 min at room temperature. After fixation, cells underwent three additional PBS washes (3 min each) and were permeabilized with 0.5% Triton X-100 for 20 min at room temperature. Following three 10-min PBS washes, cells were blocked with 5% bovine serum albumin (BSA) for 30 min at room temperature.

Cells were then incubated with primary antibody against pMLKL either overnight at 4 °C or for 2 h at room temperature. After three 10-min PBS washes, samples were incubated with fluorescent secondary antibody (Abbkine, Cat. No. A23620) for 45 min in the dark. Following final washes (three times, 10 min each with PBS), fluorescence images were acquired using a confocal microscope.

#### Live/Dead Cell Staining and Viability Assessment

RAW 264.7 murine macrophage cells were seeded in 6-well plates at a density of 1×10⁶ cells per well and cultured overnight. Cells were then treated with PBS, CBL0137, or H1N1 virus for 2 hours. Following treatments, the medium was replaced with fresh PBS, Man-NC, Glc-A4, A4, or Man-A4 solutions for 24-hour incubation. After treatment completion, cells were washed twice with PBS to remove residual compounds.

A working solution containing 2 μM Calcein-AM and 4 μM propidium iodide (PI) was prepared according to the Live/Dead Cell Double Staining Kit instructions. The staining solution was added to completely cover the cell monolayer, followed by incubation at 37 °C for 30 minutes in the dark. After incubation, cells were gently washed with PBS to remove excess dye. Viable cells (displaying green Calcein-AM fluorescence) and dead cells (displaying red PI fluorescence) were visualized and imaged using fluorescence confocal microscopy.

#### Hoechst 33342 Staining

Following experimental treatments, cells were washed three times with PBS and incubated with Hoechst 33342 staining solution (Sangon Biotech, Cat. No. E607302) diluted 1:100 in PBS at 37 °C for 30 minutes. After staining, cells were washed three times with PBS to remove excess dye, and nuclear images were acquired using fluorescence microscopy for subsequent analysis.

#### Quantitative Real-Time PCR (qRT-PCR) Analysis

Total RNA was isolated using the Universal RNA Extraction Kit (Takara, Cat. No. 9767). RNA concentration and purity were determined with a Multiscan Sky spectrophotometer (Thermo Fisher Scientific, USA). cDNA was synthesized from equal amounts of RNA using the PrimeScript 1st Strand cDNA Synthesis Kit (Takara, Cat. No. 6110A). Quantitative real-time PCR was performed with SYBR Premix (GenStar, Cat. No. A304-10) on a QuantStudio 3 system (Thermo Fisher Scientific, USA). All reactions were run in triplicate, and GAPDH served as the internal reference gene for normalization. Relative gene expression in experimental groups was calculated using the 2^(-ΔΔCT) method, with the control group set to 1.0. Primer sequences used in this study are provided in Table S4.

#### Antipneumonia Efficacy of Man-A4 in H1N1-Induced Pneumonia Mice

To assess the anti-pneumonia efficacy of mannosylated aptamer constructs, healthy male Balb/c mice (4 weeks old) were randomly divided into multiple experimental groups (n=20 per group): (1) normal control (PBS without viral infection), (2) virus control (H1N1 + PBS), (3) Man-NC treatment (H1N1 + Man-NC), (4) Glc-A4 treatment (H1N1 + Glc-A4), (5) A4 treatment (H1N1 + A4), and (6) Man-A4 treatment (H1N1 + Man-A4) with concentration of Man-A4 varies from 5 mg/kg to 30 mg/kg. All treatments were administered intranasally once daily for 4 consecutive days, beginning 24 hours post-viral inoculation.

Body weight was recorded daily and expressed as percentage change relative to initial weight (day 0). For histopathological assessment, major organs (heart, liver, spleen, lungs, kidneys, and brain) were collected at designated time points, fixed in 4% paraformaldehyde, and embedded in paraffin. Tissue sections (4 μm thickness) were subjected to hematoxylin and eosin (H&E) staining for morphological evaluation and TUNEL staining for apoptosis detection in pulmonary tissues. Survival rates were monitored throughout the study period in a separate cohort of animals.

#### Evaluation of A4 Aptamer Effects on ADAR1 RNA-Editing Activity

A dual-fluorescent reporter construct (TagBFP–mNeonGreen) containing a premature termination codon (UAG) was employed to assess A-to-I RNA editing activity. The guide RNA (LEAPER) was designed according to a previously validated strategy that recruits endogenous ADAR1 to the reporter transcript for site-specific editing^13^. Cells were co-transfected with the reporter plasmid and either LEAPER gRNA alone or LEAPER gRNA together with the A4 aptamer. After 48 h of incubation, fluorescence signals were analyzed by flow cytometry, and editing efficiency was calculated from the ratio of mNeonGreen to TagBFP fluorescence intensities. No significant difference in editing efficiency was observed between the control and A4 aptamer–treated groups.

#### Statistical Analysis

Statistical analyses were conducted using GraphPad Prism 9 and Origin 2022 software. Data are presented as mean ± s.d.. Survival data were analyzed using the Kaplan-Meier method. Statistical significance was assessed with an unpaired two-tailed Student’s t-test, one-way ANOVA, and Tukey’s multiple comparisons test. The following p-values were considered significant: ns (not significant), *P* > 0.05, **P* < 0.05, ***P* < 0.01, ****P* < 0.001.

#### Animal License

All animal procedures were conducted according to the Chinese Regulations for the Administration of Affairs Concerning Experimental Animals and all the procedures were approved by the Committee for Animal Experimentation at the Nankai University (SYXK (Jin) 2020-0007).

### Table S1. Sequence and relative abundance of the most enriched aptamer candidates.

| Family | Name | Sequence | Frequency | Total |
| --- | --- | --- | --- | --- |
| 1 | A37 | TTCAGCACTCCACGCATAGCCGACCCCGCACCGCCCTGTGTGGTCACGGCATCCCGCCTATGCGTGCTACCGTGAA | 0.29% | 3.47% |
|  | A20 | TTCAGCACTCCACGCATAGCCCGCCCACCGCCGATGTTGTTGTTGCCCTCCGCTCGCCTATGCGTGCTACCGTGAA | 1.58% |  |
|  | A22 | TTCAGCACTCCACGCATAGCGCGCACCCCCCACTCATATGGTCCGTCCGTCGCTCTCCTATGCGTGCTACCGTGAA | 1.17% |  |
|  | A32 | TTCAGCACTCCACGCATAGCGCGCGCACCCGACCCATTTGTATTGTCCTGCACCGGCCTATGCGTGCTACCGTGAA | 0.43% |  |
| 2 | A15 | TTCAGCACTCCACGCATAGCCCGGCGTGAAGACCCGTGTGCTTTCAGGTCGCCCGCCCTATGCGTGCTACCGTGAA | 3.16% | 25.60% |
|  | A39 | TTCAGCACTCCACGCATAGCCGGGCCAGACAACCAACGACCTTCTCCGTCTCGCGCCCTATGCGTGCTACCGTGAA | 0.22% |  |
|  | A24 | TTCAGCACTCCACGCATAGCCGGGCCTGCGCAACACGACTCTCTTCGGCCGCACACCCTATGCGTGCTACCGTGAA | 0.88% |  |
|  | A13 | TTCAGCACTCCACGCATAGCCCGGCCGACGGAACAATCCGTGTTCACTCGTCGCACCCTATGCGTGCTACCGTGAA | 3.80% |  |
|  | A41 | TTCAGCACTCCACGCATAGCCGGGCCACCGAGGGGGTATCATGTCCTGGTCCGCACCCTATGCGTGCTACCGTGAA | 0.18% |  |
|  | A25 | TTCAGCACTCCACGCATAGCACGCCGGACGCAACCTCCATTCCATGCGCTGCCCACCCTATGCGTGCTACCGTGAA | 0.64% |  |
|  | A17 | TTCAGCACTCCACGCATAGCTCACCCCACGCCGCGACAGTATTGCAGCCCGCCCGCCCTATGCGTGCTACCGTGAA | 1.99% |  |
|  | A30 | TTCAGCACTCCACGCATAGCACACCCCCGGCCACTGACTAAGCTTGGTCTGCTCGCCCTATGCGTGCTACCGTGAA | 0.47% |  |
|  | A12 | TTCAGCACTCCACGCATAGCACACGCGCGAAGACCCCCACCGATCCGGCCGCTCGTCCTATGCGTGCTACCGTGAA | 3.68% |  |
|  | A33 | TTCAGCACTCCACGCATAGCCCGCCGACCCTGATCTGTACGTCTTCGTCTGCCCCGCCTATGCGTGCTACCGTGAA | 0.40% |  |
|  | A36 | TTCAGCACTCCACGCATAGCCCGACCAACACCTCCGAACCGGTTTTGGCACGCCCGCCTATGCGTGCTACCGTGAA | 0.33% |  |
|  | A6 | TTCAGCACTCCACGCATAGCCCGCACCCAGCCGCGACTGCCCCCTGGTTCGGCCCGCCTATGCGTGCTACCGTGAA | 4.97% |  |
|  | A10 | TTCAGCACTCCACGCATAGCACACGACCGAACTCGCCAACCCTGTCGTCCCGCCCGCCTATGCGTGCTACCGTGAA | 4.21% |  |
|  | A28 | TTCAGCACTCCACGCATAGCTCGACCCCACACACACGACATCCTCTCTCCCGCCCGCCTATGCGTGCTACCGTGAA | 0.50% |  |
|  | A42 | TTCAGCACTCCACGCATAGCACGCCCGGACACAGATTTACTCCTCGTCCACGCCGGCCTATGCGTGCTACCGTGAA | 0.17% |  |
| 3 | A11 | TTCAGCACTCCACGCATAGCCCGGTGCCGACCGAACCACCGTTGCCCTCCGCATGCCCTATGCGTGCTACCGTGAA | 3.80% | 18.14% |
|  | A29 | TTCAGCACTCCACGCATAGCACGCCCCGAACAGGACCCCCGGCGCCCTTCCCCCGACCTATGCGTGCTACCGTGAA | 0.49% |  |
|  | A14 | TTCAGCACTCCACGCATAGCACGCCGCCGAAGCAAACCCACGTCCTCGCCCACGGCCCTATGCGTGCTACCGTGAA | 3.27% |  |
|  | A16 | TTCAGCACTCCACGCATAGCACACCGCCCCCCTCTACTCGGCACACTCGCCCGCCCCTATGCGTGCTACCGTGAA | 2.98% |  |
|  | A4 | TTCAGCACTCCACGCATAGCACGCCCGCCACCCCCTCTCCTGGCCATCCGCGCACACCTATGCGTGCTACCGTGAA | 5.67% |  |
|  | A18 | TTCAGCACTCCACGCATAGCACGCACTCCGGCCCCCCTCCAGTCCTCGCCGCACCCCCTATGCGTGCTACCGTGAA | 1.93% |  |
| 4 | A7 | TTCAGCACTCCACGCATAGCGACGGGCCAACACCCCAGGTTCGCGTCCCCGCCTGCCCTATGCGTGCTACCGTGAA | 4.79% | 38.08% |
|  | A19 | TTCAGCACTCCACGCATAGCCACGCCCGCCCAAGCTACCCTCGCCGCCGTCGGTGTCCTATGCGTGCTACCGTGAA | 1.93% |  |
|  | A27 | TTCAGCACTCCACGCATAGCCACGCGCCCTTCCACCCTAATAACACTCCGGCCCGCCCTATGCGTGCTACCGTGAA | 0.50% |  |
|  | A5 | TTCAGCACTCCACGCATAGCCCGCCCCCCGCTACTCCCCTCCTGAGTTGCCGTCGCCCTATGCGTGCTACCGTGAA | 5.61% |  |
|  | A35 | TTCAGCACTCCACGCATAGCCTGCCACGCACCTGACCCCTATTCATTTGCCCGCACCCTATGCGTGCTACCGTGAA | 0.36% |  |
|  | A23 | TTCAGCACTCCACGCATAGCCGCGCGCCACCAGATCGACACTTCGCTCGCCCCGCCCCTATGCGTGCTACCGTGAA | 0.94% |  |
|  | A9 | TTCAGCACTCCACGCATAGCCGACCACCCCCCCGACCCGGTGTCACTCCCCGCGCCCCTATGCGTGCTACCGTGAA | 4.44% |  |
|  | A2 | TTCAGCACTCCACGCATAGCCGGCCAGCCAAGTCCCCGCTCCTCACGGCCCCCCGCCCTATGCGTGCTACCGTGAA | 6.20% |  |
|  | A3 | TTCAGCACTCCACGCATAGCCGACCCGCGCCGCATTCACTGCCTACTGCCCGCCGCCCTATGCGTGCTACCGTGAA | 5.79% |  |
|  | A1 | TTCAGCACTCCACGCATAGCCGGACGCGCCCCTCTTCCTTGTGTGCTCTCGCCCGCCCTATGCGTGCTACCGTGAA | 6.55% |  |
|  | A31 | TTCAGCACTCCACGCATAGCCACGCACCCCCGCCAAACTGTACTTCCTGGCCCAGTCCTATGCGTGCTACCGTGAA | 0.44% |  |
|  | A26 | TTCAGCACTCCACGCATAGCCCGCACAACGCACGTCGACACCTGGTTCGCACCGTCCCTATGCGTGCTACCGTGAA | 0.53% |  |
| 5 | A21 | TTCAGCACTCCACGCATAGCCCGCCTAGCCCCACCCGTATACCTGTTCCGCCCCGACCTATGCGTGCTACCGTGAA | 1.23% | 6.67% |
|  | A8 | TTCAGCACTCCACGCATAGCCCGCCACCACCAATCCTCACACTAGTCCGGCCCGGTCCTATGCGTGCTACCGTGAA | 4.62% |  |
|  | A34 | TTCAGCACTCCACGCATAGCCCGCCCAGCACCGCATCACCCTGTCCACGCCCGGTCCTATGCGTGCTACCGTGAA | 0.36% |  |
|  | A38 | TTCAGCACTCCACGCATAGCCCGCCCCCCCGAACTGCCATCCGAGTCGCGCCGCTGTCCTATGCGTGCTACCGTGAA | 0.27% |  |
|  | A40 | TTCAGCACTCCACGCATAGCCCGCGACCCGACCCACCAATTGCGTCCCTCCTCGGTCCTATGCGTGCTACCGTGAA | 0.19% |  |

Note: The primer regions of the aptamer are highlighted in red.

### Table S2. Kinetic rate constants for the interaction between aptamers and ADAR1-Zα.

| Ligand - Target | k_on_ (M⁻¹s⁻¹) | k_off_ (s⁻¹) | Kinetic *K*_D_ (nM)* |
| --- | --- | --- | --- |
| A1 - ADAR1-Zα | (5.04 ± 0.63) × 10^4^ | (1.60 ± 0.15) × 10⁻^3^ | 31.7 ± 5.0 |
| A2 - ADAR1-Zα | (3.66 ± 0.88) × 10^4^ | (9.40 ± 0.12) × 10⁻^4^ | 25.7 ± 6.2 |
| A3 - ADAR1-Zα | (4.60 ± 0.90) × 10^4^ | (7.40 ± 0.08) × 10⁻^4^ | 16.1 ± 3.7 |
| A4 - ADAR1-Zα | (2.33 ± 0.39) × 10^5^ | (4.80 ± 0.05) × 10⁻^4^ | 2.1 ± 0.4 |
| A5 - ADAR1-Zα | (2.51 ± 2.04) × 10^5^ | (4.90 ± 0.07) ×10⁻^4^ | 2.0 ± 0.3 |
| A6 - ADAR1-Zα | (4.90 ± 0.39) × 10^4^ | (9.90 ± 0.07) × 10⁻^4^ | 20.2 ± 2.2 |
| A7 - ADAR1-Zα | (4.98 ± 0.31) × 10^4^ | (1.38 ± 0.13) × 10⁻^3^ | 27.7 ± 3.1 |
| A8 - ADAR1-Zα | (9.92 ± 0.22) × 10^3^ | (9.90 ± 0.10) × 10⁻^4^ | 99.8 ± 24.6 |
| A9 - ADAR1-Zα | (4.94 ± 0.84) × 10^4^ | (1.31 ± 0.16) × 10⁻^3^ | 26.5 ± 5.5 |
| A10 - ADAR1-Zα | (2.60 ± 0.50) × 10^4^ | (2.58 ± 0.43) × 10⁻^3^ | 99.2 ± 25.3 |
| A11 - ADAR1-Zα | (1.34 ± 0.07) × 10⁴ | (1.39 ± 0.11) × 10⁻^3^ | 103.7 ± 10.0 |
| A12 - ADAR1-Zα | (1 44 ± 0.35) × 10⁴ | (1.53 ± 0.19) × 10⁻³ | 104.2 ± 29.2 |
| A13 - ADAR1-Zα | (5.12 ± 1.11) × 10⁴ | (1.51 ± 0.21) × 10⁻³ | 29.5 ± 7.6 |
| A14 - ADAR1-Zα | (1.12 ± 0.01) × 10⁴ | (1.17 ± 0.07) × 10⁻³ | 96.7 ± 11.4 |
| A15 - ADAR1-Zα | (1.37 ± 0.01) × 10⁴ | (1.41 ± 0.20) × 10⁻³ | 102.9 ± 17.0 |
| A16 - ADAR1-Zα | (1.27 ± 0.01) × 10⁴ | (1.31 ± 0.10) × 10⁻³ | 103.1 ± 11.9 |

Note: Kinetic parameters were obtained from global fitting of the sensorgrams. The curves were fitted to a 1:1 Langmuir binding model using Biacore Evaluation Software. *The kinetic *K*_D_ was calculated as k_off_ / k_on_.

### Table S3. Binding kinetics of ADAR1–Zα and ZBP1-Zα with Z-DNA.

| Ligand - Target | k_on_ (M⁻¹s⁻¹) | k_off_ (s⁻¹) | Kinetic *K*_D_^#^ (nM**)** | Equilibrium *K*_D_^*^ (nM) |
| --- | --- | --- | --- | --- |
| Z-DNA - ADAR1-Zα | (3.36 ± 0.39) × 10⁵ | (1.76 ± 0.08) × 10⁻^2^ | 52.3 ± 6.3 | 44.7 |
| Z-DNA - A4-ADAR1-Zα | (5.51 ± 0.84) × 10^5^ | (3.14 ± 0.05) × 10⁻^3^ | 5.7 ± 0.9 | 4.7 |
| Z-DNA - ZBP1-Zα | (1.59 ±0.15) × 10^5^ | (3.34 ± 0.06) × 10⁻^2^ | 210.0 ± 15.9 | 254.9 |

Note. ^#^The equilibrium *K*_D_ value was derived from a direct fit of the concentration-dependent binding curve at equilibrium. *The kinetic *K*_D_ was calculated as k_off_ / k_on_.

### Table S4. Primer sequences used for qPCR.

| **Name** | **Forward** | **Reverse** |
| --- | --- | --- |
| GAPDH (mouse) | ACAGTCAGCCGCATCTTCTT | ACGACCAAATCCGTTGACTC |
| IL-18 (mouse) | AAAGTGCCAGTGAACCC | TTTGATGTAAGTTAGTGAGAGTGA |
| IL-1β (mouse) | AGCTACGAATCTCCGACCAC | CGTTATCCCATGTGTCGAAGAA |
| IL-6 (mouse) | AAAGAGGCACTGGCAGAAAA | TTTCACCAGGCAAGTCTCCT |
| TNF-α (mouse) | CAGAAAGCATGATCCGCGACG | CAGTAGACAGAAGAGCGTGGT |
| IFN-β (mouse) | AGCTCCAAGAAAGGACGAACA | GCCCTGTAGGTGAGGTTGAT |
| GAPDH (human) | GTCTCCTCTGACTTCAACAGCG | ACCACCCTGTTGCTGTAGCCAA |
| IL-18 (human) | GATAGCCAGCCTAGAGGTATGG | CCTTGATGTTATCAGGAGGATTCA |
| IL-1β (human) | CCACAGACCTTCCAGGAGAATG | GTGCAGTTCAGTGATCGTACAGG |
| IL-6 (human) | AGACAGCCACTCACCTCTTCAG | TTCTGCCAGTGCCTCTTTGCTG |
| TNF-α (human) | CTCTTCTGCCTGCTGCACTTTG | ATGGGCTACAGGCTTGTCACTC |
| IFN-β (human) | CTTGGATTCCTACAAAGAAGCAGC | TCCTCCTTCTGGAACTGCTGCA |

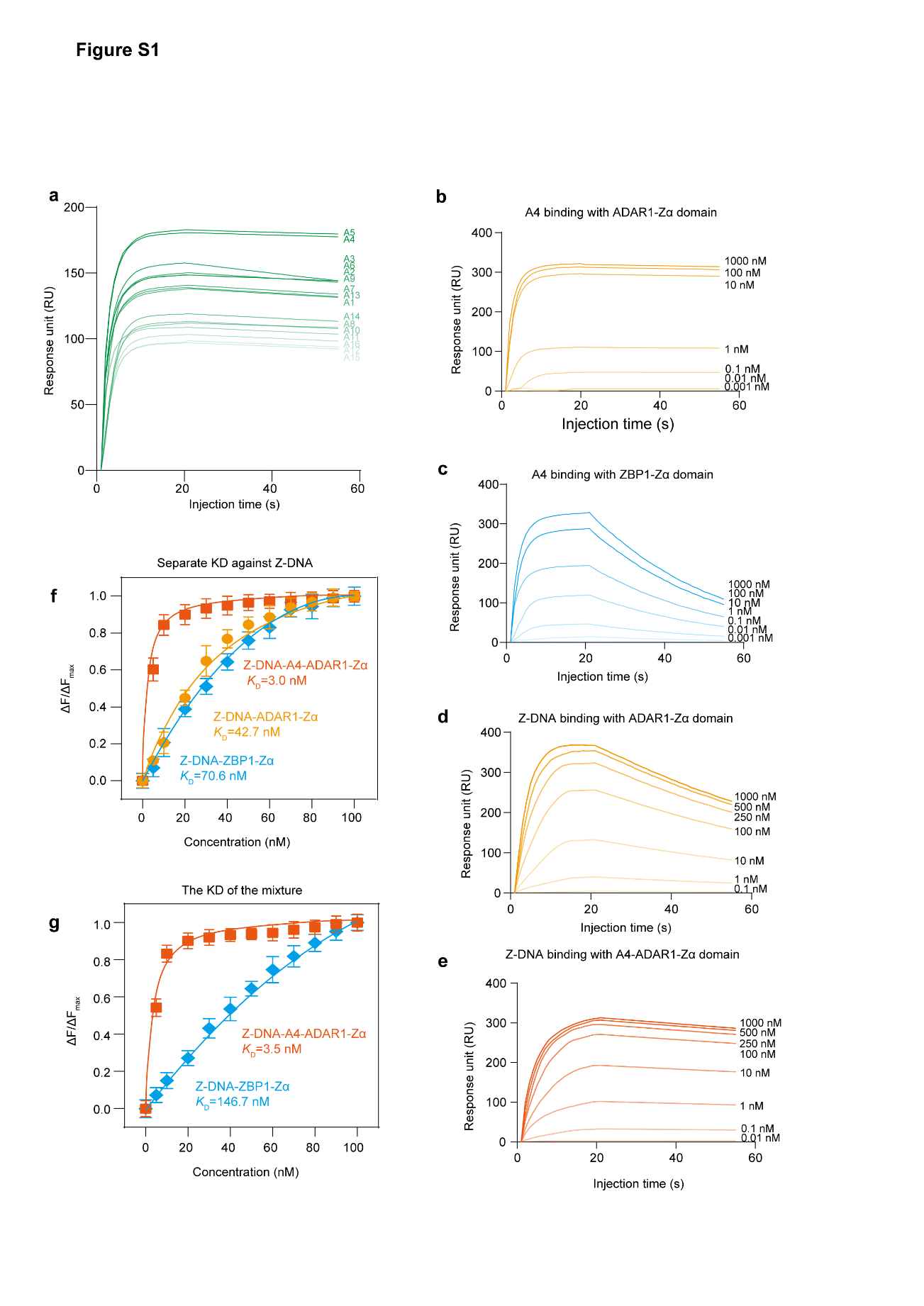

### Figure S1. Aptamer screening and validation of A4 binding properties using SPR and HTCFQ assays. (a) SPR sensorgrams for the top 16 candidate aptamers binding to ADAR1-Zα. (b-d) Concentration-dependent SPR analysis for binding affinity determination: (b) A4 binding to ADAR1-Zα, (c) A4 binding to ZBP1-Zα, (d) Z-DNA binding to ADAR1-Zα, and (e) Z-DNA binding to the pre-formed A4-ADAR1-Zα complex. (f) Binding curves and equilibrium dissociation constants (*K*_D_) for ADAR1-Zα, A4-ADAR1-Zα complex, and ZBP1-Zα binding to Z-DNA, as determined by HTCFQ. (g) Competitive binding curves and *K*_D_ values for a mixture of A4-ADAR1-Zα and ZBP1-Zα against Z-DNA, measured by HTCFQ. Data are presented as mean ± s.d. (n = 3).

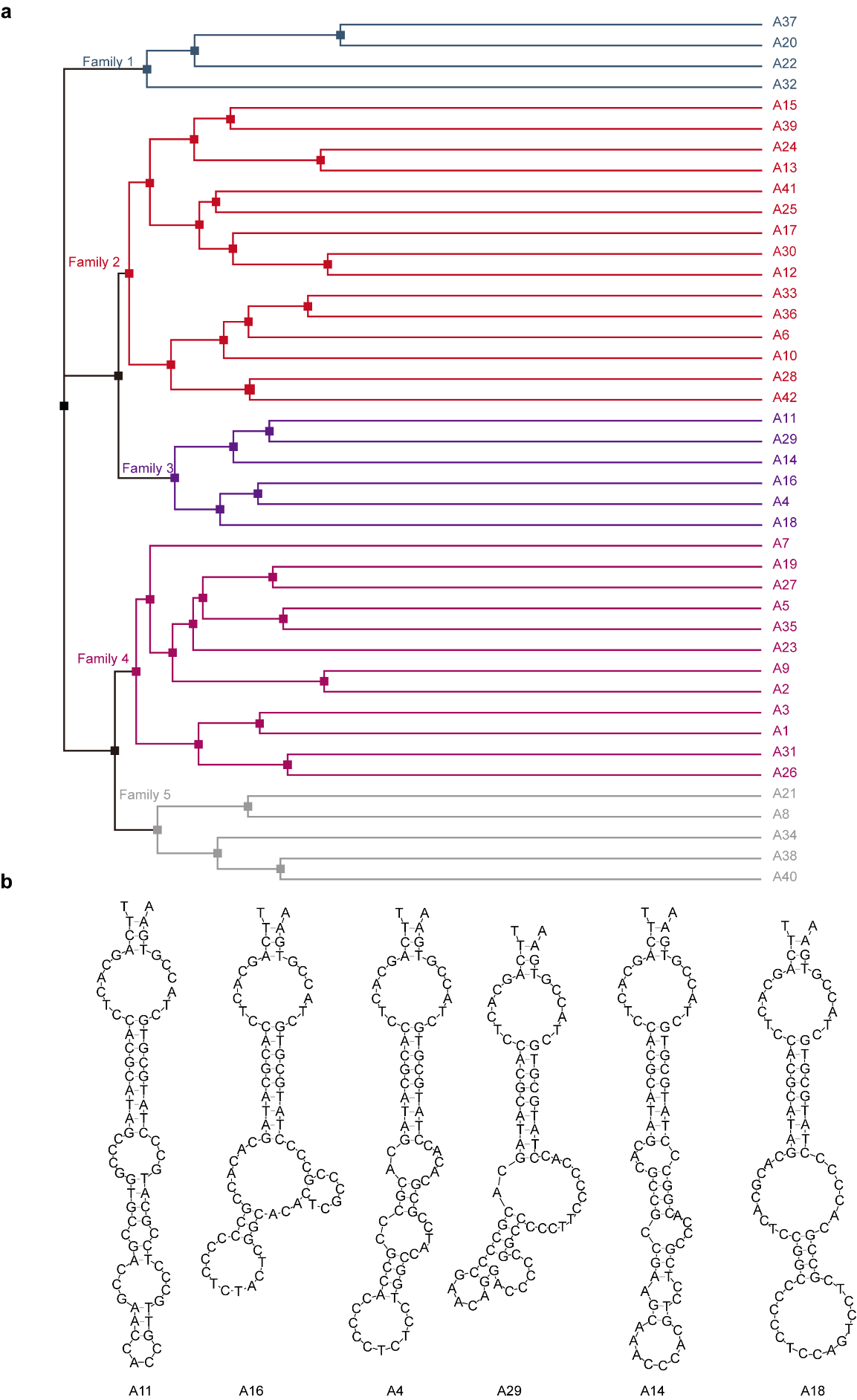

### Figure S2. (a) Phylogenetic trees of similarity among 42 aptamers constructed using the Clustal omega. (b) Secondary structures of aptamers from Family III.

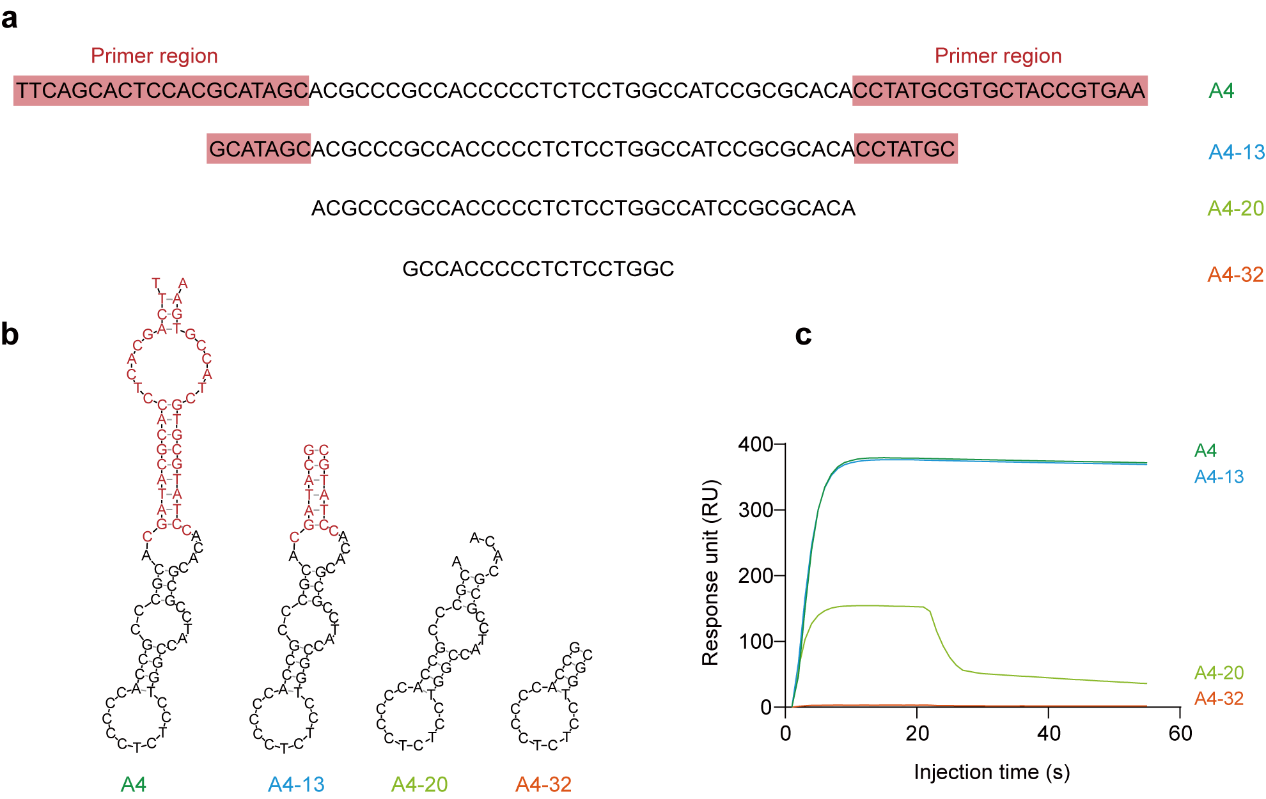

### Figure S3. Characterization of the full-length A4 aptamer and it truncates. (a) Sequences of the full-length A4 aptamer and three truncates (A4-13, A4-20, A4-32). The predicted core binding regions are highlighted in color. (b) Predicted secondary structures of the aptamers from (a). (c) Representative SPR sensorgrams demonstrating the binding of the aptamers to ADAR1-Zα. The nomenclature (A4-13, A4-20, A4-32) refers to the number of bases truncated from one end of the full-length A4.

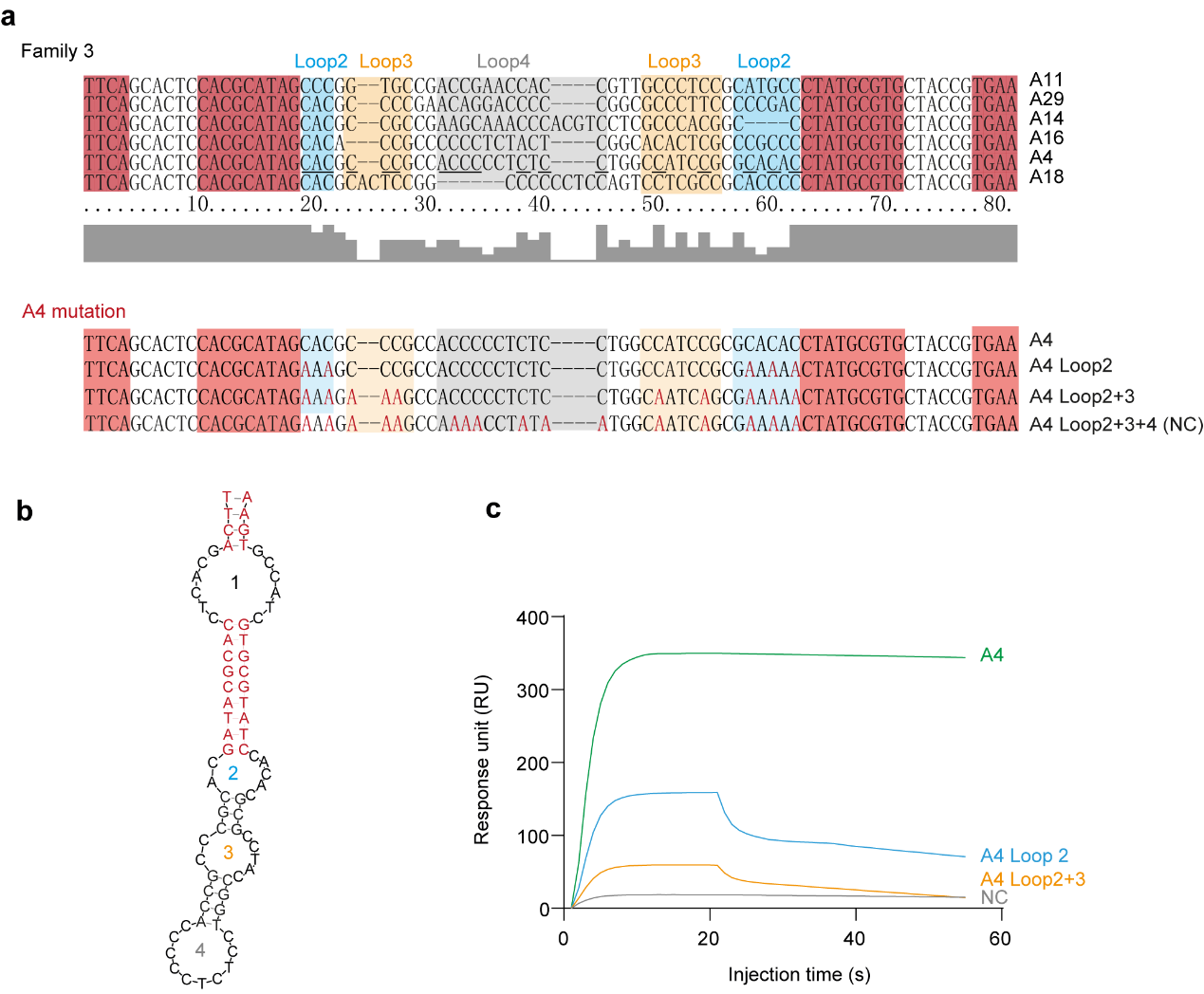

### Figure S4. Characterization of the A4 loop mutant variants. (a) Sequence alignment of aptamers belonging to Family III. Nucleotides conserved with A4 are underlined. Residues highlighted in red correspond to the stem regions illustrated in (b). (b) Predicted secondary structure of A4, with loop regions corresponding to those labeled in the sequence alignment (a) numerically indicated. (c) SPR sensorgrams comparing the binding of A4 loop mutants to ADAR1-Zα.

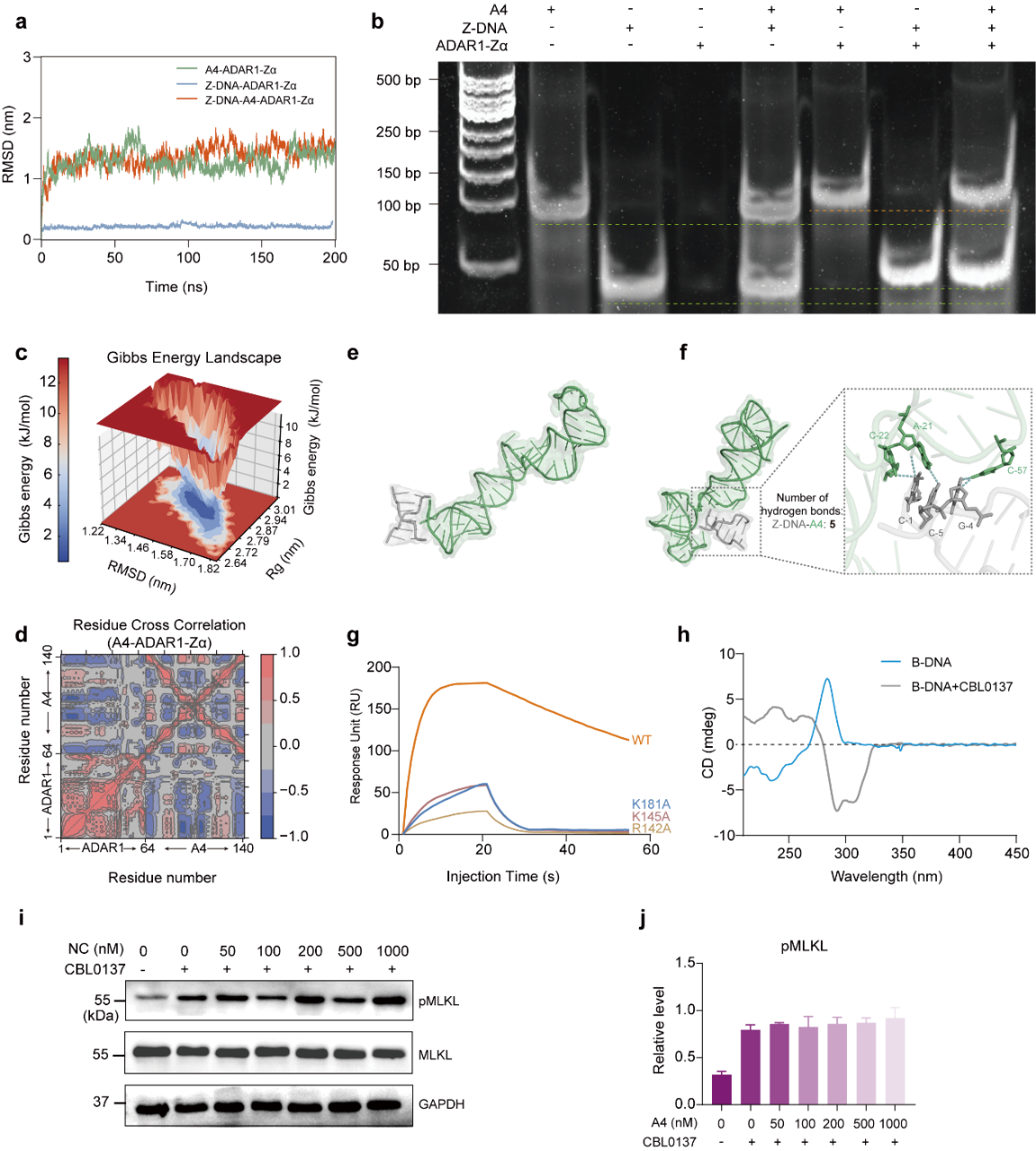

### Figure S5. Binding analysis of A4 to ADAR1-Zα. (a) RMSD plot from molecular dynamics simulations comparing A4-ADAR1-Zα, Z-DNA-ADAR1-Zα and Z-DNA-A4-ADAR1-Zα complexes. (b) SPS-PAGE analysis providing direct evidence for ternary complex formation, as indicated by a distinct band shift only when ADAR1-Zα, Z-DNA, and the A4 aptamer are all present. (c) Gibbs energy landscape from molecular dynamics simulation showing the stable conformational state of A4 when bound to ADAR1-Zα. Rg represents the radius of gyration, while RMSD stands for root-mean-square deviation. (d) Residue cross-correlation analysis showing significant dynamic interaction between ADAR1-Zα domain (1-64 residues) and A4 (65-140 residues). (e) The binding pattern of Z-DNA to A4 without allostery. The region of A4 that Z-DNA binds with does not belong to A4’s target binding region. (f) A close-up of the hydrogen binding interactions of Z-DNA to A4 after the A4 allostery. (g) SPR binding curves of WT and mutated ADAR1-Zα against Z-DNA. (h) Verification of the CBL0137-induced B- to Z-DNA transition by CD spectroscopy. (i) Western blot analysis confirming that the A4 Loop2+3+4 mutant, serving as a negative control (NC), does not reduce pMLKL levels in A549 cells after 24-hour treatment across increasing concentrations. (j) Quantitative analysis of pMLKL expression from (i), demonstrating the absence of dose-dependent effects by the mutant control.

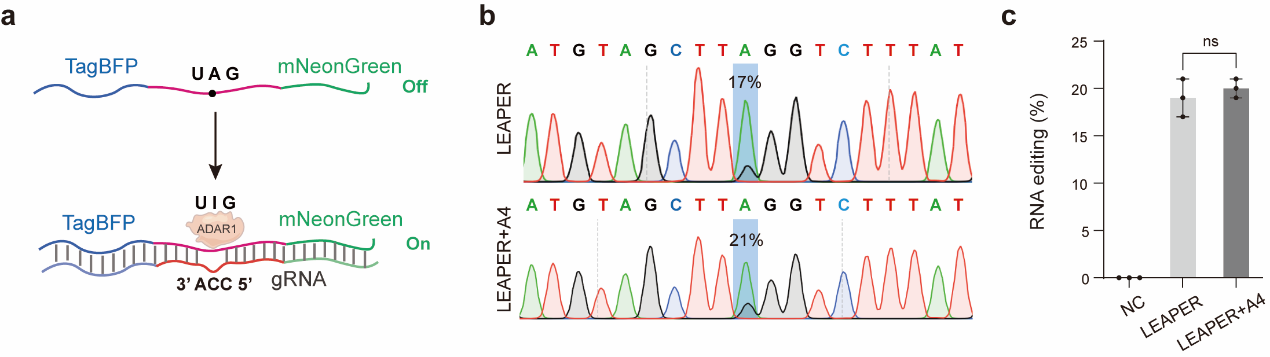

### Figure S6. The A4 aptamer does not alter the endogenous RNA-editing activity of ADAR1. (a) Schematic of the dual-fluorescent reporter system (TagBFP‑mNeonGreen) used to monitor A-to-I editing efficiency. The LEAPER gRNA directs ADAR1 to a specific site on the reporter transcript. (b) Representative Sanger sequencing chromatograms from reporter transcripts under the indicated conditions, showing no change in the editing level. Target sites are highlighted in blue regions. (c) Quantification of editing efficiency from replicate experiments (n = 3). Data are presented as mean ± s.d.; differences are not statistically significant (ns, p > 0.05 by one-way analysis of variance (ANOVA) test).

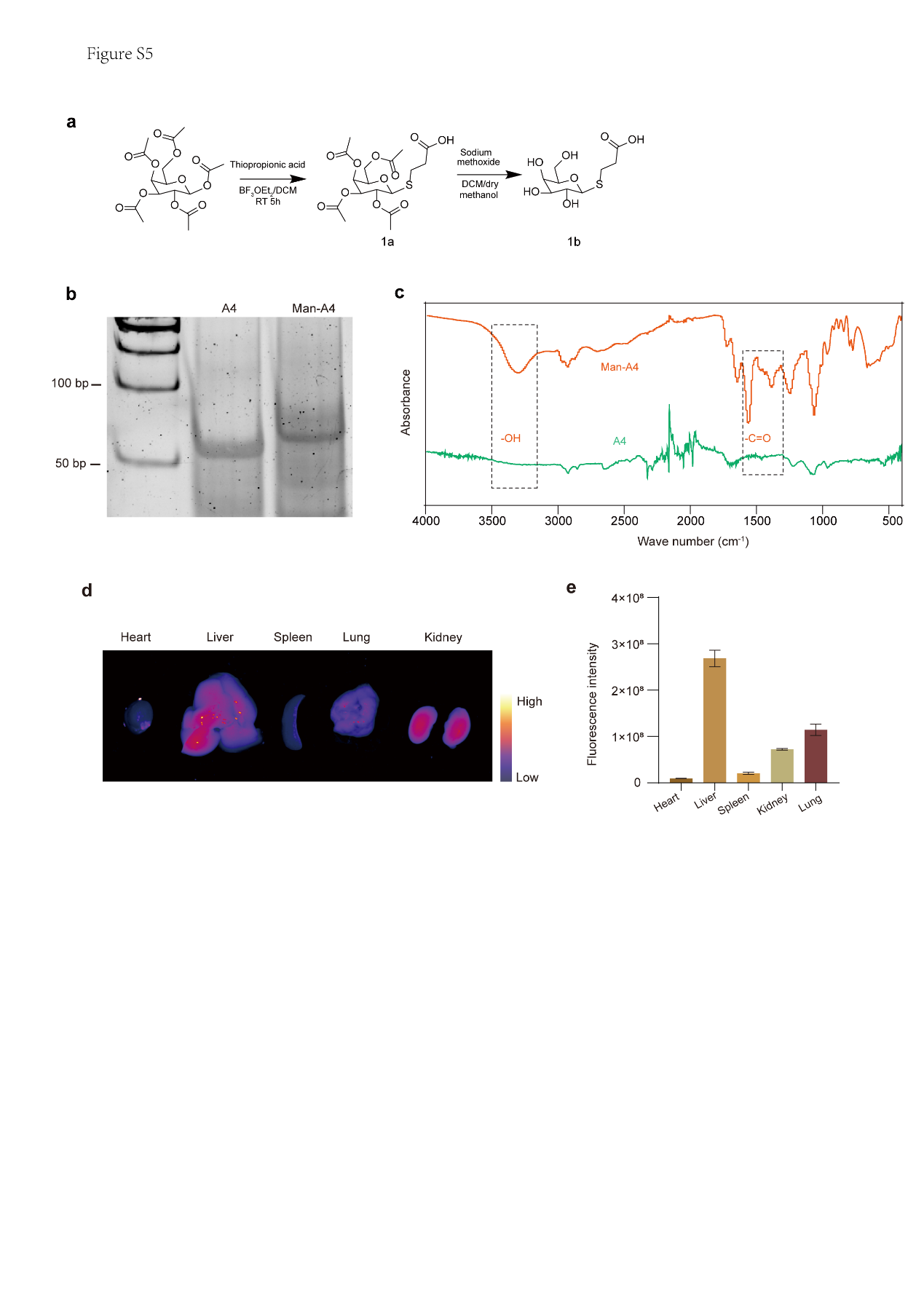

### Figure S7. Synthesis and characterization of mannosylated A4 (Man-A4) aptamer. (a) Schematic illustration of the synthetic route for 3-hydroxybut-3-en-1-yl-thio-mannoside (1b). (b) Polyacrylamide gel electrophoresis (PAGE) analysis comparing the migration of unmodified A4 and Man-A4. (c) Infrared spectroscopy analysis of A4 (10 μM) and Man-A4 (30 μM) for chemical structure validation. (d) Organ distribution of Man-A4-Cy5 at 48 hours post intratracheal administration. (e) Quantification of the fluorescence intensity shown in (d).

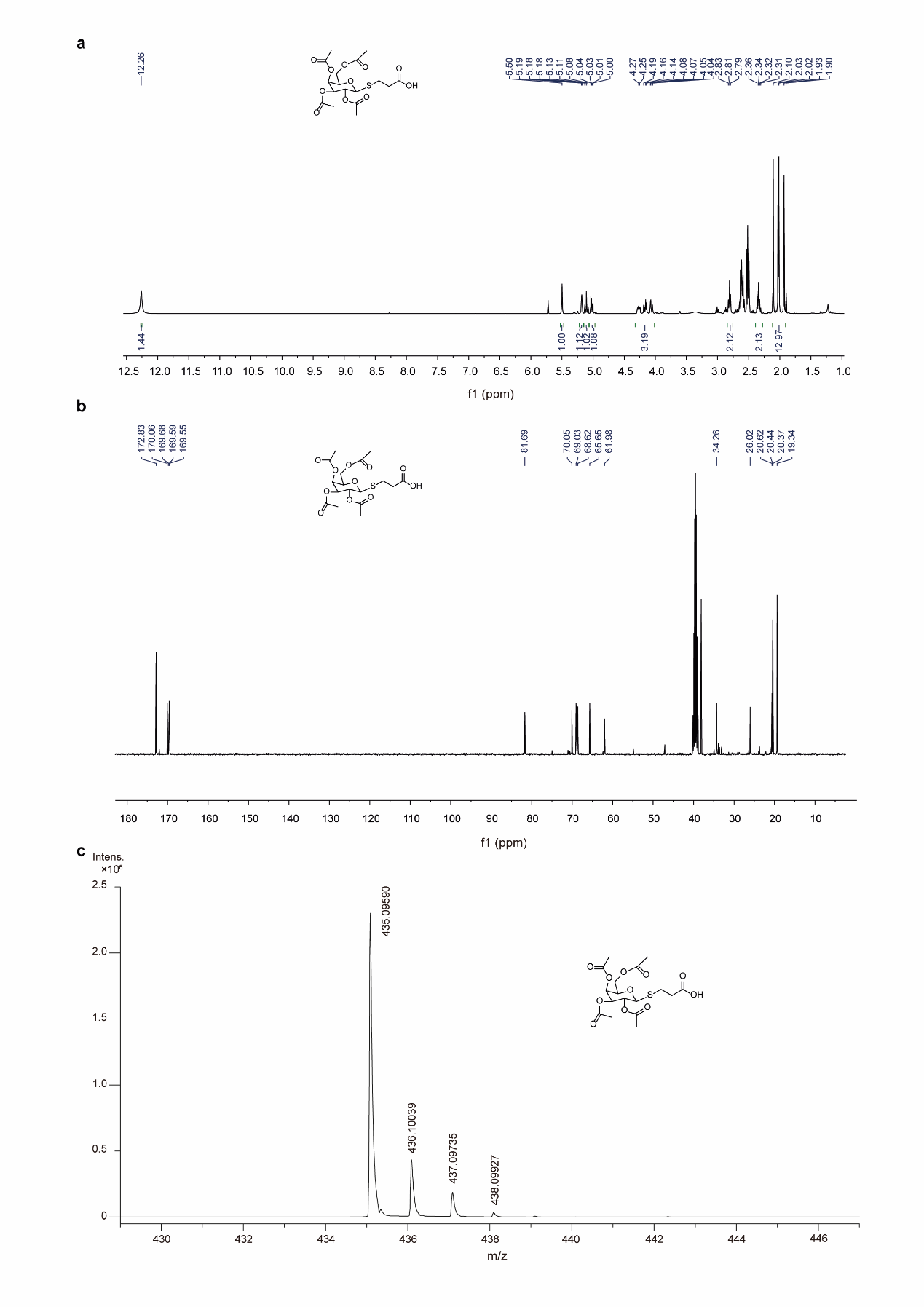

### Figure S8. ^1^H NMR spectrum (a), ^13^C NMR spectrum (b) and high-resolution mass spectrum (c) of 1a shown in Figure S7a.

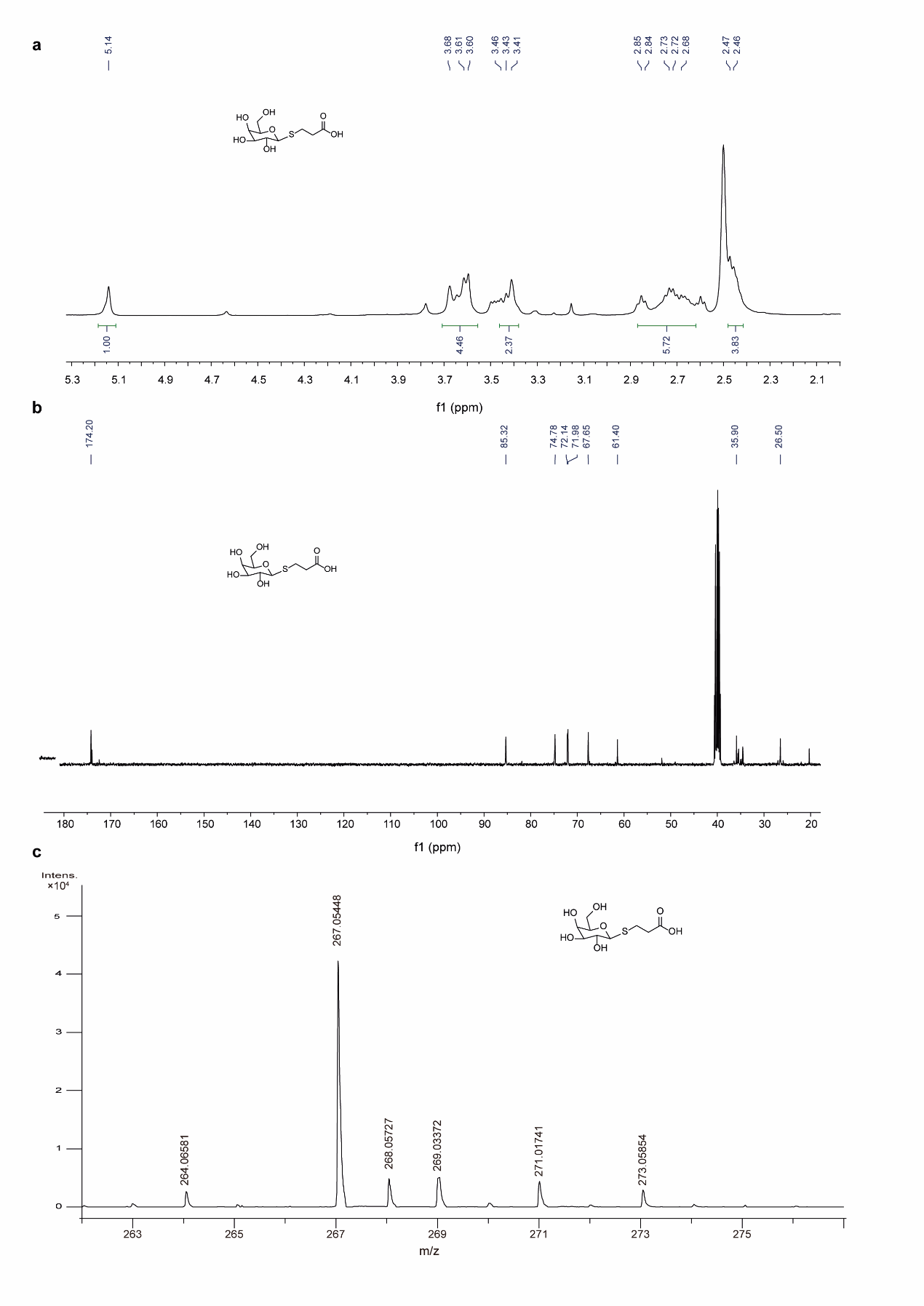

### Figure S9. ^1^H NMR spectrum (a), ^13^C NMR spectrum (b) and high-resolution mass spectrum (c) of 1b shown in Figure S7a.

.

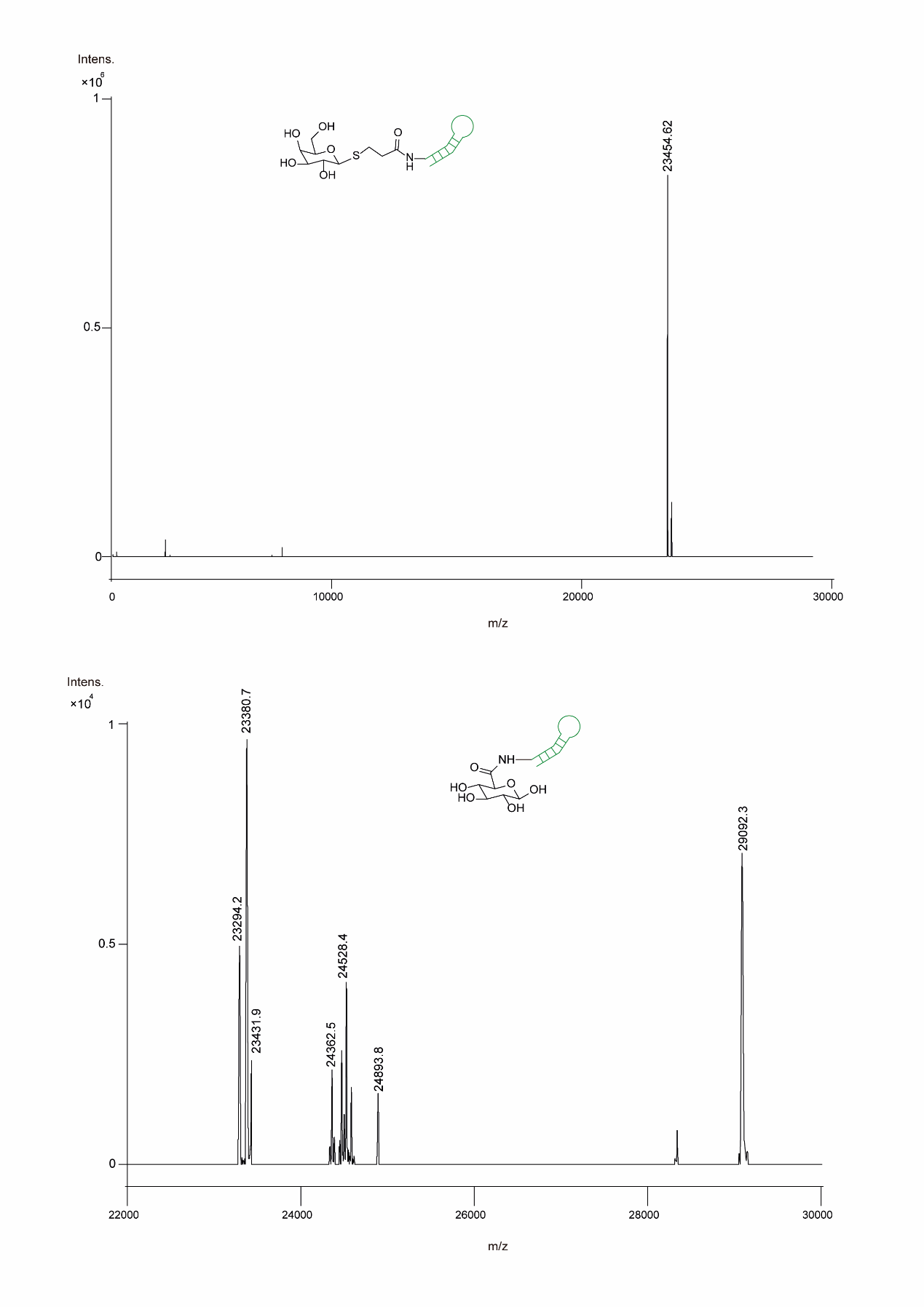

### Figure S10. The high-resolution mass spectrum of Man-A4 (top) and Glc-A4 (bottom) shown in Figure S7a.

.

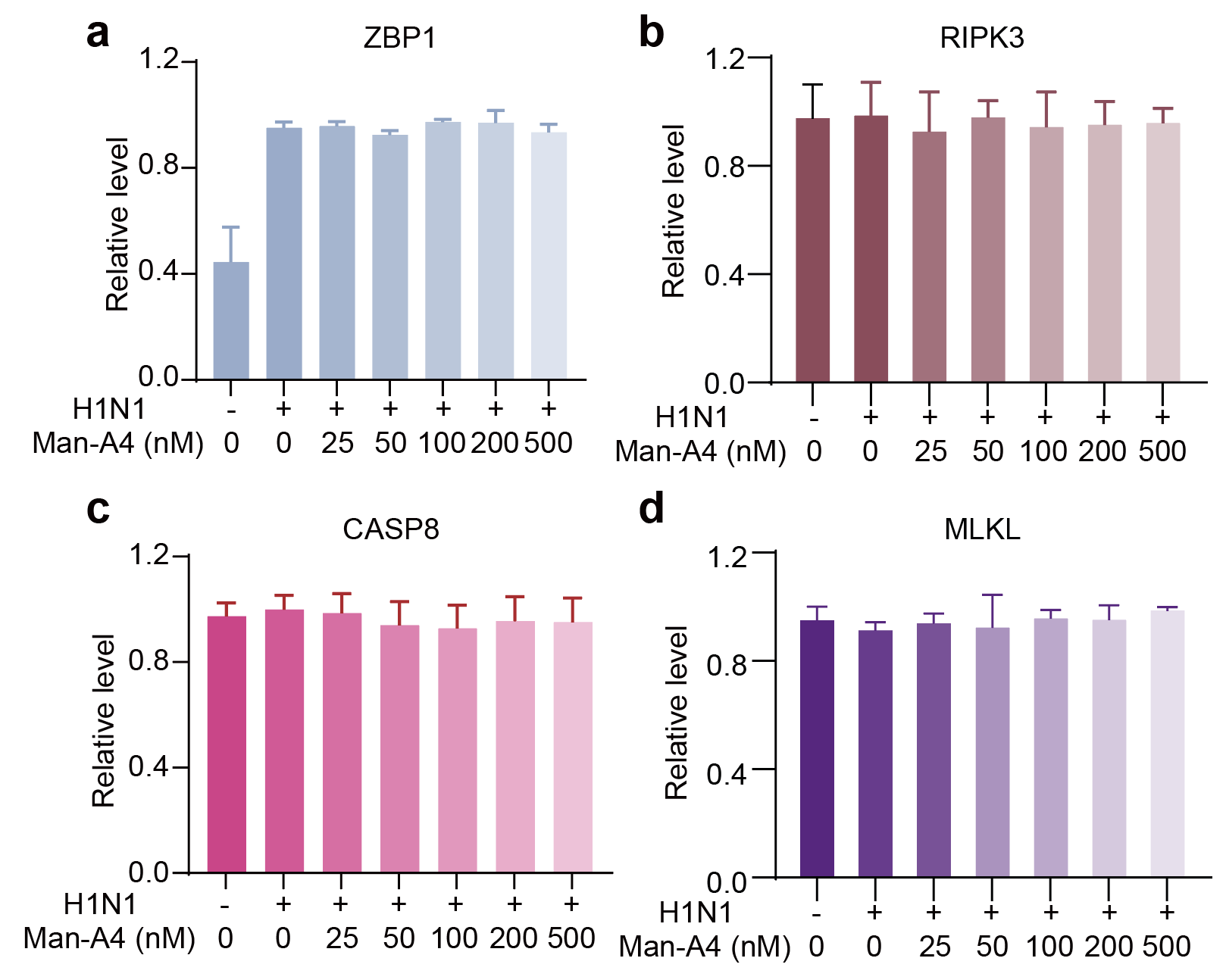

### Figure S11. Quantification of (a) ZBP1, (b) RIPK3, (c) CASP8, and (d) MLKL protein expression from the Western blots in Figure 4a. Data are shown as mean ± s.d. (n = 3 independent experiments).

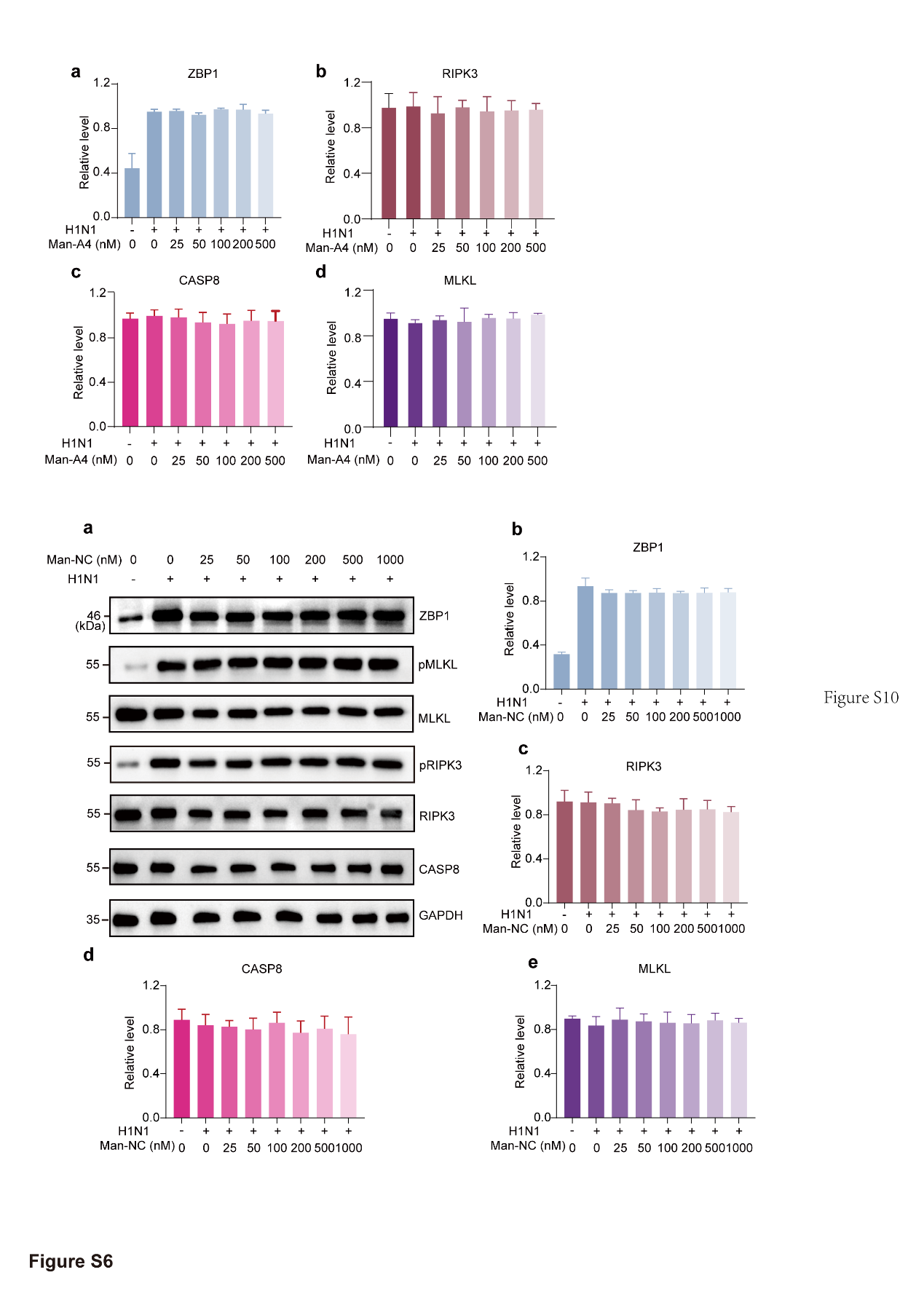

### Figure S12. Evaluation of Man-NC effects on necroptosis pathway components in RAW 264.7 cells. (a) Western blot analysis of key necroptosis-related proteins (ZBP1, MLKL, pMLKL, RIPK3, pRIPK3, Caspase-8) in RAW 264.7 cells treated with increasing concentrations of Man-NC. (b-e) Quantitative analysis of (b) ZBP1, (c) RIPK3, (d) Caspase-8, and (e) MLKL protein expression levels from (a). Data are presented as mean ± s.d. from three independent experiments.

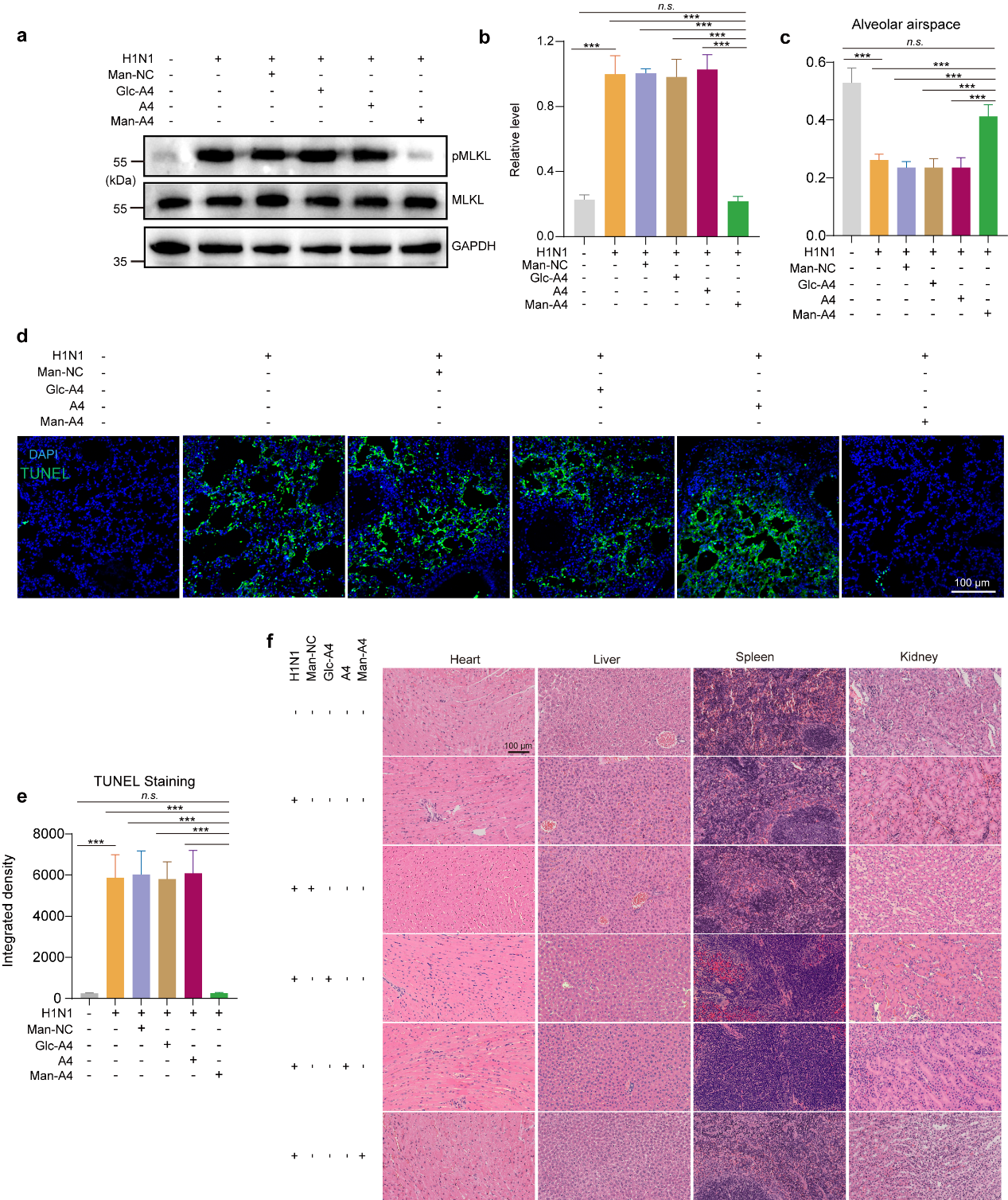

### Figure S13. Therapeutic efficacy and safety evaluation of Man-A4 *in vivo*. (a) Western blot analysis of pMLKL and total MLKL expression in lung tissues from mice treated with H1N1, Man-NC, Glc-A4, or Man-A4. (b) Quantification of the pMLKL/MLKL ratio from (a), showing significant suppression of necroptosis signaling by Man-A4. (c) Morphometric analysis of alveolar airspace area in lung tissues across treatment groups. (d) Representative TUNEL staining images of lung sections, indicating DNA fragmentation as a marker of cell death. (e) Quantitative analysis of TUNEL-positive cells from (d), demonstrating reduced apoptosis in the Man-A4-treated group. (f) H&E-stained sections of heart, liver, spleen, and kidney tissues, confirming the biosafety of Man-A4. All data are presented as mean ± s. d. (n = 3 biologically independent samples). Statistical significance was determined by Student’s *t*-test: **p < 0.01, ***p < 0.001.
